# Preventing vision loss in a mouse model of Leber Congenital Amaurosis by engineered tRNA

**DOI:** 10.1101/2025.07.10.660754

**Authors:** Enes Akyuz, Pawan K. Shahi, Lionel Gissot, Ahmad Al Saneh, Divya Sinha, Giovanni M. Hanstad, Meha Kabra, Maria A. Fernandez Zepeda, David M. Gamm, Samuel M. Young, Christopher A Ahern, Bikash R. Pattnaik

**Author notes:** **Correspondence:** Bikash R. Pattnaik, 1300 University Avenue, SMI 112, Madison, WI, 53706. **EA** and **PKS** are co-first authors and contributed equally.

## Abstract

Premature termination codons (PTCs) are associated with rare genetic disorders. Inducing targeted read-through of these ‘nonsense mutations’ presents a potential therapeutic strategy for modifying disease outcomes. We previously reported that one such PTC, W53X, in the *KCNJ13* gene causes blindness and Leber congenital amaurosis type-16 (LCA-16) due to loss of function of the inwardly rectifying potassium channel 7.1 (Kir7.1). Here, we present the proof of concept of a therapeutic approach based on anticodon-engineered transfer RNA (ACE-tRNA). The ACE-tRNA encodes the amino acid tryptophan (Trp) and suppresses the W53X PTC, restoring full-length protein expression. We used helper-dependent adenovirus (HDAd) to deliver the ACE-tRNA^Trp.UAG^ (tRNA^Trp.UAG^) and rescue Kir7.1 function and physiology in patient-specific human induced pluripotent stem cell-derived retinal pigment epithelium (hiPSC-RPE) cells. Furthermore, in a W53X mouse model of LCA16, HDAd delivery of tRNA^Trp.UAG^ resulted in durable restoration of vision as measured by retinography. This study provides the first example of the therapeutic application of ACE-tRNA for treating an inherited form of blindness.

## Introduction

Nonsense mutations introduce premature termination codons (PTCs) into mRNA, leading to truncated, non-functional proteins that can cause severe genetic diseases^1^. Nonsense mutations account for approximately 15% of all inherited disorders, including cystic fibrosis, Duchenne muscular dystrophy, and certain forms of congenital blindness^2^. In these diseases, PTCs prevent the production of full-length functional proteins, which often result in progressive and devastating symptoms. Innovative therapeutic strategies are needed for PTCs to produce functional proteins.

Several strategies to overcome nonsense mutations have been tested, including pharmacological read-through agents, aminoglycoside antibiotics, and gene-editing techniques^3^. Both read-through drugs and aminoglycoside antibiotics, such as gentamicin, can induce partial read-through of PTCs by allowing near-cognate tRNA incorporation^4^. However, these agents frequently cause unpredictable amino acid substitutions, which can lead to proteins with compromised functions if missense variants are poorly tolerated^5^. Gene editing by CRISPR/Cas9 precisely corrects mutations at the DNA level, but it is mutation-specific, requires efficient but transient expression of the editor, and extensive testing of potential off-target editing, especially in post-mitotic cells such as those in the retina^6-8^. More relevant for PTC therapies is that many nonsense mutations cause ‘n of 1’ ultra-rare diseases and require a positional and ideally gene-agnostic approach that provides specific encoding of the correct amino acid in the PTC.

Anticodon-engineered transfer RNAs (ACE-tRNAs) suppress nonsense mutations by using the translation machinery to incorporate the correct amino acids at premature stop codons^9-11^. This approach bypasses the need for genomic alterations and avoids nonspecific encoding associated with small-molecule read-through agents^12^. ACE-tRNAs have been previously shown to rescue a variety of PTC-containing mRNAs *in vitro*, including CFTR^13,14^, CDKL5^15^, and the cardiac potassium channel HERG^16^. However, no known cases exist of successfully rescuing a PTC-bearing gene *in vivo* to achieve a therapeutic outcome. Notably, AAV-mediated tRNA delivery for Hurler Syndrome results in poor rescue *in vivo*^17^ although such viral approaches could be helpful for tRNA delivery^18^.

We previously reported that Leber Congenital Amaurosis type 16 (LCA16), a form of early-onset retinal dystrophy, is caused by a nonsense mutation, c.158G>A (p.W53X), in the *KCNJ13* gene (NM_002242.4)^20^. *KCNJ13* encodes inwardly rectifying potassium channel 7.1 (Kir7.1), which is essential for maintaining retinal pigment epithelium (RPE) cell function and retinal ionic homeostasis^19^. The nonsense mutation results in a loss-of-function mutant of the Kir7.1 channel, leading to progressive photoreceptor degeneration and vision loss. Our primary goal was to develop a novel therapy for LCA16 that can selectively target RPE cells within the retina^20^.

Here, we present a proof-of-concept study of ACE-tRNA, specifically engineered to recognize the tryptophan nonsense codon (ACE-tRNA^Trp. UAG^) to restore Kir7.1 W53X mutant channel function in cellular and animal models of LCA16. In induced pluripotent stem cell-derived RPE (hiPSC-RPE) from an LCA16 patient (LCA16 hiPSC-RPE), we determined that helper-dependent adenovirus (HDAd)-delivered ACE-tRNA^Trp. UAG^ (tRNA^Trp.UAG^) led to a read-through of the W53X mutation, resulting in a full-length functional Kir7.1 protein localized to the cell membrane. Subretinal delivery of the HDAd ACE-tRNA^Trp.UAG^ (HD tRNA^Trp.UAG^) in a *Kcnj13*-mutant mouse model restored RPE cell function and vision, as evidenced by improved electroretinogram (ERG) responses. Together, these results demonstrate that therapeutic ACE-tRNA holds promise as a targeted treatment for LCA16 and other genetic disorders caused by nonsense mutations.

## Results

### Suppression of UAG Premature Termination Codons by ACE-tRNA^**Trp.UAG**^

To assess UAG nonsense codon read-through by codon-edited tryptophan tRNA, we tested if anticodon-modified tRNA rescued the expression of a super-folder GFP harboring an in-frame TAG mutation at N150 (sfGFP^TAG^). This site has been previously shown to tolerate a wide variety of side-chain sizes and thus can help predict TAG suppression and stop codon read-through^21^. In the human genome, GtRNAdb (https://gtrnadb.ucsc.edu) and tryptophan tRNA (tRNA^Trp^) are encoded by seven iso-decoder genes (TRW-CCA 1-1 through 5-1) with a CCA anticodon. We screened six CCA iso-decoders (TRW-CCA 2-1 through 5-1).

In humans, the tRNA tryptophan gene encodes a tRNA specific to tryptophan (tRNA-Trp) that decodes the UGG codon in the mRNA. This tRNA’s anticodon CCA base pairs with the UGG codon during translation^22^. We screened seven CCA iso-decoder genes (TRW-CCA 2-1 through 5-1) engineered to carry CUA anticodons for their ability to read through the in-frame UAG codon of sfGFP^TAG^ reporter protein. We performed fluorescence imaging and flow cytometry analysis in HEK293T cells transfected with the sfGFP^TAG^ construct to assess the translation read-through efficiency (**Fig. 1A**). Co-expression of engineered tRNA-Trp-CUA based on tRNA-Trp-CCA-2-1 (hereafter termed **tRNA**^**Trp.UAG**^) led to >42% of GFP-positive cells 24 h post-transfection (**Fig. 1B, H**). All other engineered Trp tRNA isodecoders were less efficient (tRNA-Trp-CCA-3-1 (33.5% GFP expression), tRNA-Trp-CCA-4-1 (41.7%), and tRNA-Trp-CCA-5-1 (29.9%), **Fig. 1C-E, H**). Although all tRNA-Trp-CCAs had AUC in the tRNA anticodon loop structure, the nucleotide sequence shows few mismatches within the tRNA structural loop domains that may be important for ribosomal binding and could influence read-through efficiency (**Fig. 1B-E)**. In control experiments, cells with the pUC19 plasmid and the sfGFP^TAG^ plasmid showed no GFP expression (**Fig. 1F-H**). Overall, these data show that tRNA^Trp.UAG^ can read through the UAG stop codon in HEK293T cells, producing full-length functional GFP.

**Fig. 1:**
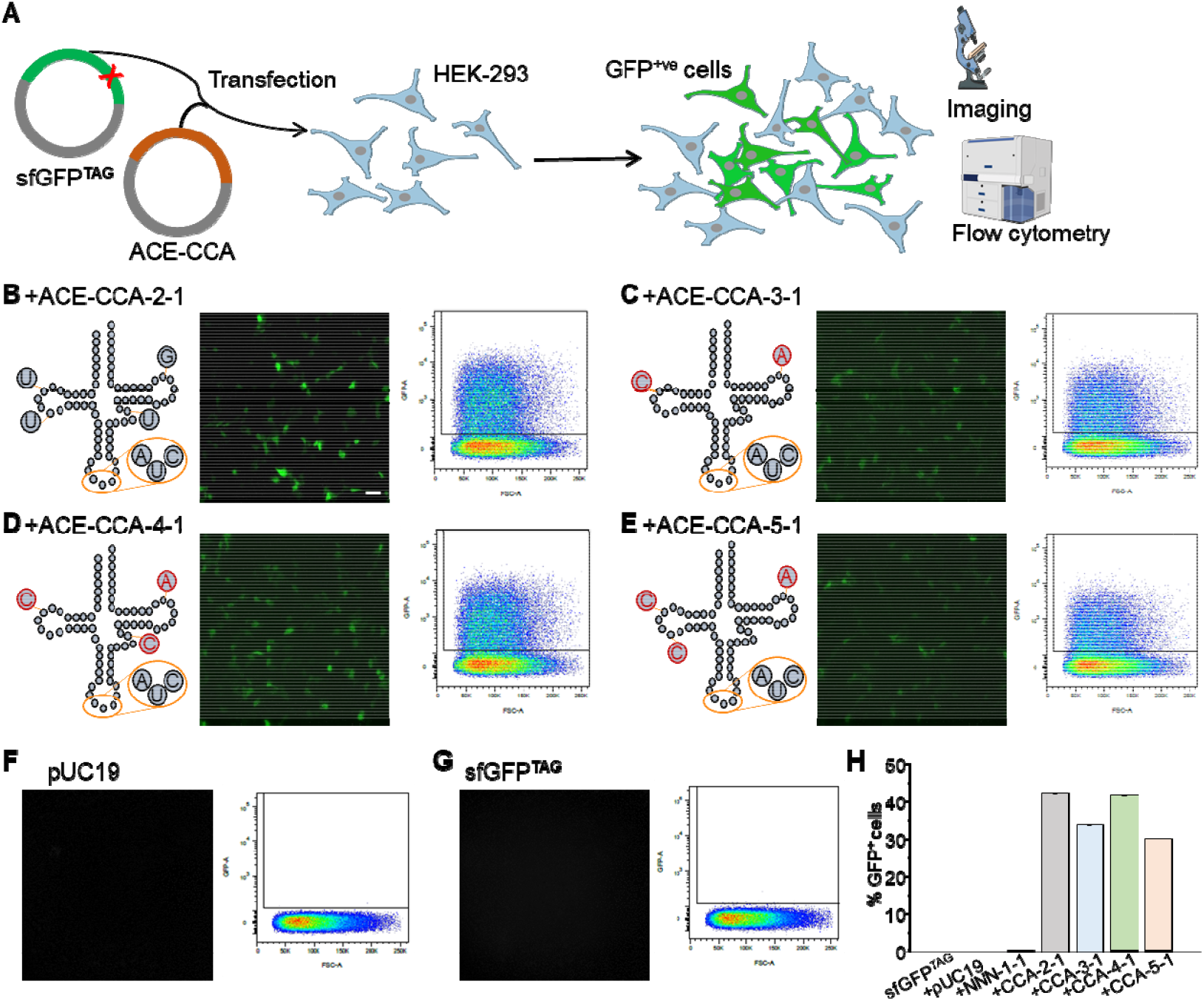
Selection of tRNA^Trp.UAG^-tRNA isodecoders using sfGFP^TAG^ readthrough. **A**, Overview of the experiments in HEK293 cells transfected with sfGFP^TAG^ (green plasmid) and various codon-edited ACE-CCA-tRNA (red plasmid) to assay GFP read-through efficiency by fluorescence imaging and flow cytometry. Co-transfection of sfGFP^TAG^ along with the codon-edited tRNAs **B**, ACE-CCA-2-1, **C**, ACE-CCA-3-1/3-2/3-3, **D**, ACE-CCA-4-1, **E**, and ACE-CCA-5-1 show representative structures and varied amounts of GFP fluorescence expression in HEK293T cells along with flow cytometry plots. Cells transfected with **F**, pUC19, or **G** sfGFP^TAG^ alone showed no GFP expression. **H**, Percentage comparison of the different isodecoders for tryptophan tRNA (ACE-CCA) to read through the GFP^TAG^ by flow cytometry assay. ACE-CCA stands for engineered tRNA isodecoder type tRNA-Trp-CCA (2-1 through 5-1). Scale bar=50 μm.

### tRNA^Trp.UAG^ leads to readthrough of exogenous Kir7.1^W53X^ and proper channel localization

To test if tRNA^Trp.UAG^ can mediate read through of an mRNA encoding ectopic *KCNJ13* with the 158G>A mutation, we initially determined the copy number of 4X tRNA^Trp.UAG^ was optimal ***(Supplemental Fig. 1)*** as it was tested earlier^14^. We then transfected HEK293 cells with plasmids encoding the heterologous N-terminally GFP-tagged Kir7.1 W53X (GFP-Kir7.1^W53X^) and tRNA^Trp.UAG^ at a 1:3 ratio to find readthrough (**Fig. 2A)**. Notably, we observed fluorescence-positive cells as evidence of biological read-through of the mutant Kir7.1 channel by tRNA^Trp.UAG^ ***(Supplemental Fig. 2)***. After 24-48 hours of transfection, cells treated with tRNA^Trp.UAG^ showed GFP fluorescence at the plasma membrane, suggesting successful trafficking of the full-length GFP-Kir7.1 channel protein. In contrast, the truncated GFP-Kir7.1^W53X^ polypeptide remained cytoplasmic without tRNA^Trp.UAG^ (**Fig. 2B**). To verify the expression of a full-length protein, we also probed the Kir7.1 C-terminus by immunocytochemistry using an anti-Kir7.1 antibody (C12). We used an anti-Na-K-ATPase antibody as a membrane marker. While cells expressing wild-type (WT) GFP-Kir7.1 showed membrane localization of the full-length protein **(Fig. 2B**, left panels), GFP-Kir7.1^W53X^ cells showed cytoplasmic localization of the tagged protein (**Fig. 2B**, middle panel). In contrast, GFP-Kir7.1^W53X^ cells co-expressing tRNA^Trp.UAG^ produced a full-length protein localized at the membrane (**Fig. 2B**, right panel). Pearson correlation coefficient analysis **(Fig. 2C**, n=22 for each group) confirmed membrane localization of GFP-Kir7.1^W53X^ in tRNA^Trp.UAG^-expressing cells compared to that in Kir7.1^W53X^ cells (P= 1.05E-15; one-way ANOVA test). Thus, tRNA^Trp.UAG^-mediated readthrough produced a full-length Kir7.1 protein trafficked to the membrane, which is necessary for channel function.

**Fig. 2:**
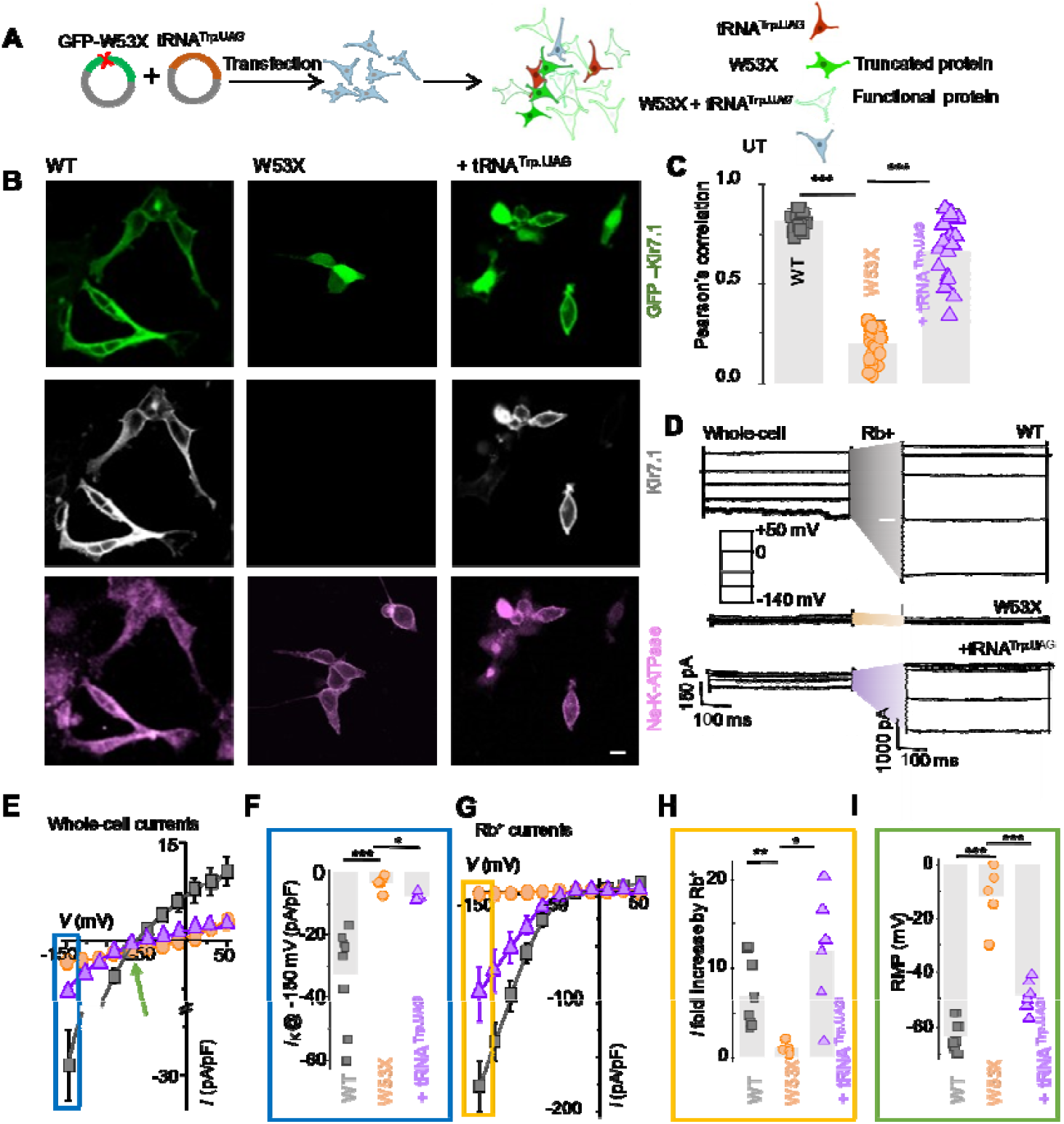
Suppression of ectopic *KCNJ13* disease mutation W53X by ACE-tRNA^Trp. UAG^. **A**. Schematic representation of W53X and ACE-tRNA^Trp. UAG^ (tRNA^Trp.UAG^) plasmids were co-transfected into HEK293 cells, which resulted in the expression of any combination of tRNA^Trp.UAG^, truncated 53 amino acid polypeptide, and full-length Kir7.1 protein channel. Cells showing positive tdTomato reporter expression represent successful tRNA^Trp.UAG^ gene expression, as shown in Supplemental Fig. 2. **B**, Confocal images show the expression of GFP fluorescence at the cell membrane due to proper localization of GFP-fused WT Kir7.1 in the cytoplasm due to truncated GFP-W53X polypeptide and a mix of membrane and cytoplasmic locations due to tRNA^Trp.UAG^ readthrough GFP-Kir7.1 protein. Both the middle and lower panels are immunocytochemistry images of cells (same as shown in the upper panel), probed with a C-terminal anti-Kir7.1 antibody (Gray) that detects only full-length protein that colocalizes with Na-K-ATPase (magenta). Scale bars in B, 20 μm. **C**, Membrane colocalization of Kir7.1 and Na-K-ATPase was measured by calculating Pearson’s coefficient for WT (black quadrangles), W53X (orange circles), and W53X + ACE-tRNA^Trp. UAG^ (purple triangles). An identical selection area was used on the cell membrane for the ROI to ensure consistency across measurements. Data points in **C** are individual ROIs that were compared for statistical significance using two-tailed Student’s *t*-test; n > 3 biological replicates. Statistical significance was determined as *\*\*\**P*<0*.*001*. **D**, Representative whole-cell current traces for cells measured in the presence of physiological K^+^ or high concentrations of Rb^+^ in the extracellular solution. Current responses were generated by applying 50 mV voltage steps (−150 to +50 mV, see inset) for 500 ms from a holding potential of 0 mV for WT, W53X alone, or W53X co-transfected with tRNA^Trp.UAG^ cells. The vertical scale bars indicate 150 pA for the K^+^ current and 500 pA for the Rb^+^ currents, whereas the horizontal scale bar is 100 ms. **E**, Average I-V plot (*I*, current density) for the Kir current measured in physiological K^+^ in WT (black quadrangles), W53X mutant (orange circles), and W53X + tRNA^Trp.UAG^ (purple triangles) cells. The blue rectangle represents the inward current density in F, and the green arrow shows the resting membrane potential in I. **F**, Comparison of K^+^ current densities for all recorded cells, as in E, measured at -150 mV. **G**, Current density plot for Rb^+^ ions recorded in cells as in E. Orange rectangle was included for the measured current increase in H. **H**, Rb^+^ enhances the inward current at -150 mV, which is represented as a fold-increase. **I**, Plot of resting membrane potentials for individual cells as in E. Data are individual recordings from n>3 biological repeats. Significance was determined as *\**P*<0*.*05, \*\**P*<0*.*01*, and ***P<0.001 using one way ANOVA. WT, GFP-Kir7.1; W53X, GFP-Kir7.1^W53X^; +tRNA^Trp.UAG^, GFP-Kir7.1^W53X^ + tRNA^Trp.UAG^.

### Restoration of Kir7.1 channel function

To test if the full-length Kir7.1 protein was functional, we co-transfected HEK293 cells with GFP-Kir7.1^W53X^ and tRNA^Trp.UAG^, as described above, tested whole-cell currents 48 h after transfection. Representative Kir7.1 channel current response to 50 mV voltage steps ranging from +50 to -150 mV over 600 ms from a holding potential of 0 mV are shown in **Fig. 2D**. We observed distinct inwardly rectifying potassium current, a hallmark of Kir7.1 channel function, in cells expressing WT Kir7.1 and in cells expressing GFP-Kir7.1^W53X^ and tRNA^Trp.UAG^, but not in cells expressing GFP-Kir7.1^W53X^ alone (**Fig. 2D**, left panels). As shown in **Fig. 2E**, WT Kir7.1 (black squares) and GFP-Kir7.1^W53X^ cells co-expressing tRNA^Trp.UAG^ (purple triangle) showed an inwardly rectifying current-voltage (I-V) curve with a large inward current for hyperpolarized potentials compared to a small outward current for depolarized potentials. The mutant GFP-Kir7.1^W53X^ channel (orange circle) showed non-rectifying linear I-V with severely reduced inward and outward currents. tRNA^Trp.UAG^-expressing GFP-Kir7.1^W53X^ cells (n=6) showed I_K_ current densities of -7.7 ± 0.4 pA/pF at -150 mV, compared to -28.4 ± 5.5 pA/pF in cells expressing the WT channel (n=10,). This was significantly improved compared to mutant channel-expressing cells (−3.3 ± 1.1 pA/pF, n=5, P=0.004; one-way ANOVA test) (**Fig. 2F**).

Similarly, in the presence of extracellular Rb^+^, an established activator of the Kir7.1 channel, the inward currents in cells expressing WT Kir7.1 and in cells expressing GFP-Kir7.1^W53X^ together with tRNA^Trp.UAG^ selectively increased, not in cells expressing GFP-Kir7.1^W53X^ alone (**Fig. 2D**, right panel). It enhanced the inward current and observed an increase in inward current density from -1.9 ± 0.1 pA/pF in GFP-Kir7.1^W53X^ expressing cells (**Fig. 2G**, orange circle) to -93.1 ± 24.9 pA/pF in cells co-expressing tRNA^Trp.UAG^ (**Fig. 2G**, purple triangles). The increased Rb^+^ current density indicates a functional Kir7.1 channel, as observed for the WT channels (−177 ± 23.1 pA/pF) (**Fig. 2G**, black squares). As shown in **Fig. 2H**, the GFP-Kir7.1^W53X^ and tRNA^Trp.UAG^ co-expressing cells exhibited an Rb^+^ fold change, measured as I_Rb/IK_ at −150 mV, of 11.7 ± 2.7, which was comparable to that of the WT channel-expressing cells (9.0 ± 2.6), more than 10-fold higher than that of the mutant channel-expressing cells (0.9 ± 0.3) (P=0.013; one-way ANOVA test). Similarly, the measured resting potential in **Fig. 2I** showed that cells co-expressing GFP-Kir7.1^W53X^ and tRNA^Trp.UAG^ showed hyperpolarization (−48.7 ± 2.3 mV). This was comparable to cells expressing the WT (−61.6 ± 2.2 mV) but significantly hyperpolarized compared to mutant channel-expressing cells (−12 ± 5.1 mV) (**Fig. 2I**, P=5.15E-07; one-way ANOVA test). The rescue of the Kir7.1 current profile, Rb^+^-induced inward current increase, and membrane potential confirmed that tRNA^Trp.UAG^ leads to the expression of a fully functional Kir7.1 channel, a critical requirement for a beneficial therapeutic outcome.

### tRNA^Trp.UAG^-mediated restoration of Kir7.1 function in patient-derived hiPSC-RPE cells

For tRNA^Trp.UAG^ to be an effective therapeutic agent, it must function efficiently within the physiological target cells. Kir7.1 channels are expressed in the apical processes in the polarized RPE cells. We, therefore, used LCA16 patient-derived hiPSC-RPE cells (which carry the *KCNJ13* W53X mutation). We transduced mature monolayer cultures with helper-dependent adenovirus (HDAd) carrying 1X (***Supplemental Fig. 3***) or 4X tRNA^Trp.UAG^ (HD tRNA^Trp.UAG^) **(Fig. 3A)**. In addition to detecting the presence of the HDAd viral genome in transduced cells by real-time PCR (***Supplemental Fig. 4***), we analyzed expression of endogenous Kir7.1 and protein localization two weeks post-transduction (**Fig. 3B, *Supplemental Fig. 5***). Immunocytochemistry with the C-terminal Kir7.1 antibody revealed that unlike LCA16 hiPSC-RPE cells (**Fig. 3B**, middle panel), WT hiPSC-RPE (**Fig. 3B**, left panel), and tRNA^Trp.UAG^-treated cells (**Fig. 3B**, right panel) exhibited Kir7.1 protein on the membrane, as the Pearson’s coefficient showed co-localization between Kir7.1 and Na-K-ATPase (N=36 for each group) **(Fig. 3C)**. Although membrane localization of Kir7.1 is of paramount importance, we did notice sub-cellular localization of readthrough proteins. Traces for whole-cell and Rb^+^ enhanced currents showed large inward currents for WT and cells transduced with HD tRNATrp.UAG compared to untreated control cells **(Fig. 3D)**. A comparison of the average whole-cell current density (**Fig. 3E**, I_K_) measured at -150 mV for WT iPSC-RPE, untreated, and treated LCA16 iPSC-RPE cells showed improvement after transduction with HD tRNA^Trp.UAG^ (P =0.0319). In addition, HD tRNA^Trp.UAG^ treatment increased the Rb^+^-induced inward current to 9.01 ± 1.78, a roughly 7-fold increase in current magnitude compared to LCA16 hiPSC-RPE control cells (1.3 ± 0.2) and comparable to WT iPSC-RPE (6.8 ± 0.6) (**Fig. 3F**, P=0.0225; one-way ANOVA test). The average resting membrane potential **(Fig. 3G)** of HD tRNA^Trp.UAG^ treated LCA16 iPSC-RPE cells was –30.5 ± 7.2 mV, comparable to–37.9 ± 2.9 mV for WT iPSC-RPE, but hyperpolarized compared to LCA16 hiPSC-RPE control cells (–4.8 ± 1.7 mV) (**Fig. 3G**, P=0.034; one-way ANOVA test). Collectively, these results demonstrate that tRNA^Trp.UAG^ can restore endogenous Kir7.1 protein, its localization to the plasma membrane, and its physiological function in patient-derived iPSC-RPE cells, supporting its therapeutic potential of tRNA-mediated nonsense read-through.

**Fig. 3:**
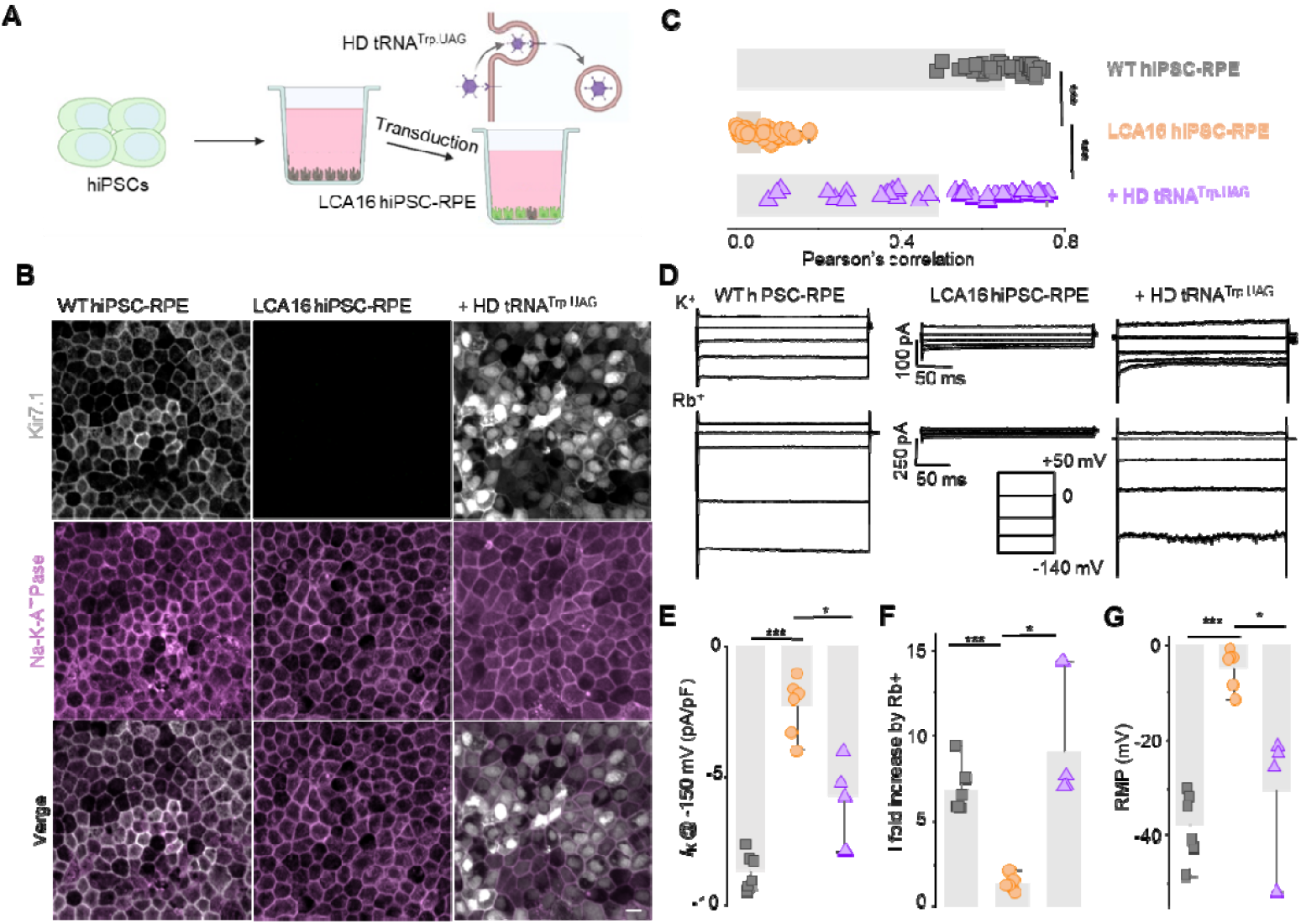
ACE-tRNA^Trp.UAG^ mediated readthrough of endogenous Kir7.1 W53X in LCA16 hiPSC-RPE cells. **A**, Schematic of LCA16 hiPSC-RPE grown to a mature tight monolayer of pigmented cells transduced with HDAd virus-packaged ACE-tRNA^Trp. UAG^ (HD tRNA^Trp.UAG^). **B**, Confocal images of WT hiPSC-RPE, LCA16 hiPSC-RPE, and LCA16 hiPSC-RPE transduced with HD tRNA^Trp.UAG^ shows the immunofluorescence localization of Kir7.1 (gray) and Na-K-ATPase (magenta). Scale bar, 20 μm. **C**, Quantitative fluorescence colocalization of Kir7.1 and Na-K-ATPase proteins by Pearson’s correlation. The co-localization of membrane and membrane proteins was analyzed by ROI selection using confocal microscopy. As shown in Fig. 3C, an identical area on the membrane was selected for the ROI, ensuring consistency across the measurements (N = 36 cells). **D**, Representative current traces in the presence of physiological K^+^ or high Rb^+^ extracellular solutions generated in response to 50 mV voltage steps from -150 to 50 mV from a holding potential of 0 mV (inset) for WT hiPSC-RPE WT, LCA16 hiPSC-RPE, and LCA16 hiPSC-RPE cells treated with HD tRNA^Trp.UAG^. The vertical scale bars indicate 100 pA for K^+^ currents and 250 pA for Rb^+^ currents, whereas the horizontal scale bar indicates 50 ms. **E**, K^+^ current densities were measured at -150 mV in individual WT hiPSC-RPE (black quadrangles), LCA16 hiPSC-RPE (orange circles), and LCA16 hiPSC-RPE cells treated with HD tRNA^Trp.UAG^ (purple triangles) groups. **F**, Inward current fold increase by extracellular Rb^+^ measured at -150 mV, and **G**, resting membrane potentials for individual cells, as in E. The data corresponds to individual cells in more than three experimental repeats, and significance was measured as*\**P*<0*.*05, \*\**P*<0*.*01*, and *\*\*\**P*<0*.*001* when LCA16 hiPSC-RPE was compared with the WT hiPSC-RPE or HD tRNA^Trp.UAG^ treated cells using one-way ANOVA.

### *In vivo* rescue of RPE cell function

To translate our observations from patient-derived hiPSC-RPE to an *in vivo* setting, we evaluated the efficacy of HD tRNA^Trp.UAG^ in a Kir7.1 W53X mouse model of LCA16^22^. First, we assessed CMV-sfGFP transduction using HDAd to determine whether this viral delivery method could transduce retinal cells *in vivo*. We observed broad GFP expression in flat-mount RPE images 14 days after subretinal HDAd virus-mediated transduction of RPE cells. This demonstrates that HDAd is competent for tRNA delivery in this model (**Fig. 4A, B**). We then administered tRNA^Trp.UAG^ through HDAd viral transduction (HD tRNA^Trp.UAG^) together with CMV-sfGFP with a TAG nonsense mutation (HD-sfGFP^UAG^) to WT mice. We observed the expression of the GFP protein in the RPE floret, indicating the ability of tRNA^Trp.UAG^ to read through the nonsense mutation in the gene and to produce a full-length protein (**Fig. 4B**). A control group injected with the HD-sfGFP^UAG^ virus alone exhibited no GFP expression, confirming that the read-through we observed was mediated by tRNA^Trp.UAG^.

**Fig. 4:**
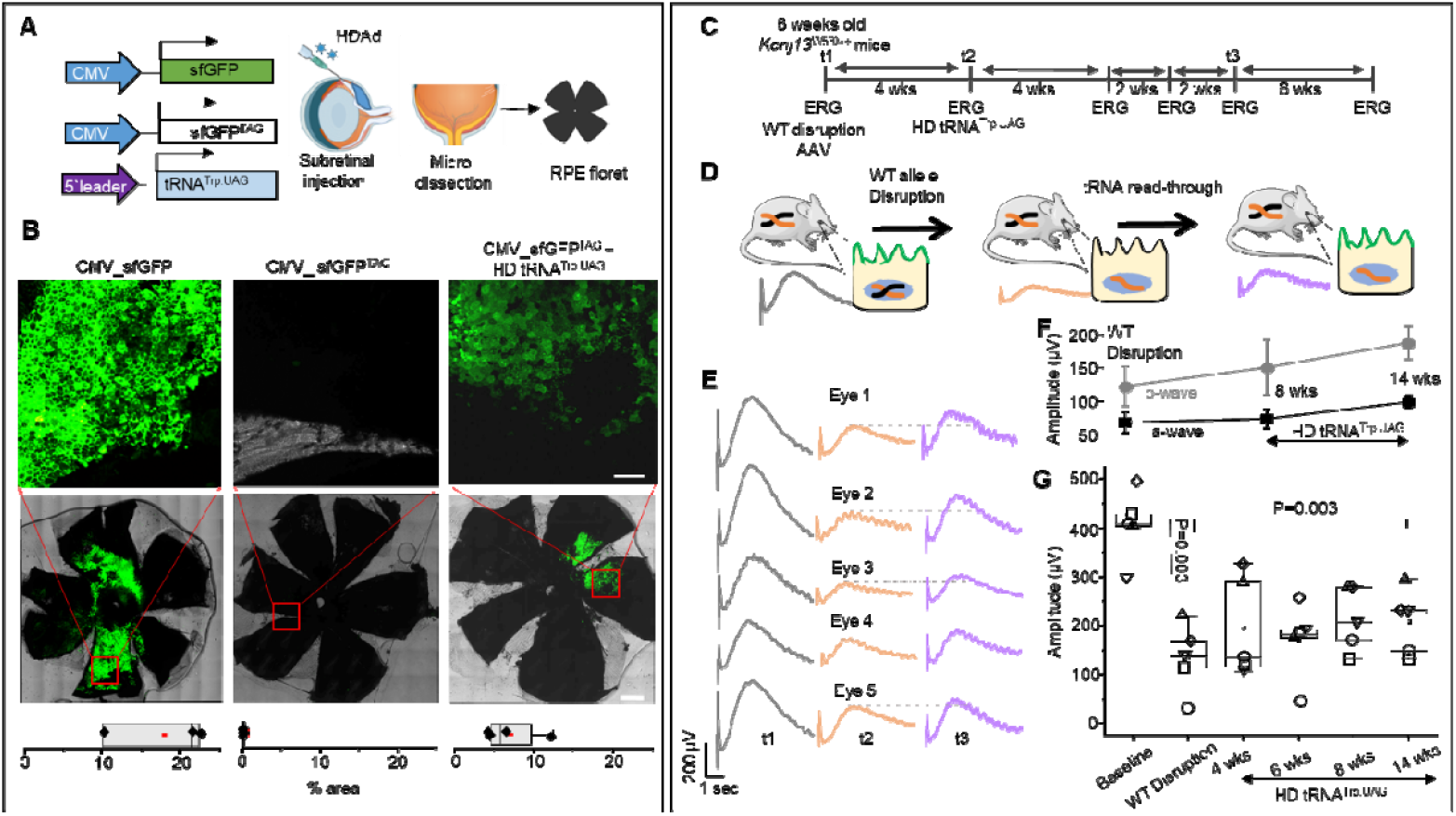
Readthrough of nonsense mutation in *Kcnj13*^W53X/ΔR^ mouse by HDAd mediated tRNA^Trp.UAG^ delivery. **A**, Schematic of HDAd virus-mediated delivery of sfGFP, sfGFP^TAG^, or sfGFP^TAG^ with 4X tRNA^Trp.UAG^ to the mouse eye via the subretinal route and RPE floret imaging. **B**, Representative RPE floret images showing fluorescence expression in mice injected with sfGFP, sfGFP^TAG^, and sfGFP^TAG^ plus 4X tRNA^Trp.UAG^. The upper panels show magnified views of the indicated areas, the middle panel images show whole florets, and the lower panel shows the percentage of fluorescence area coverage due to transduction. Scale bars=50 µm for 20X image (top) and 500 µm for the floret images (middle). **C**, Time course of the *in vivo* experiment: The WT allele of *Kcnj13*^W53X/+^ was disrupted after the baseline ERG (t1), followed by the delivery of the HDAd virus carrying therapeutic 4X tRNA^Trp.UAG^ (t2). Follow-up ERG was performed on mice at different intervals until week 14 (t3). **D**, *In vivo* experimental demonstration of RPE genotype-phenotype correlations in *the Kcnj13*^*W53X/*Δ*R*^ *mouse model* used for 4X tRNA^Trp.UAG^-mediated nonsense readthrough. **E**, The ERG c-wave comparison in five individual eyes at baseline (*Kcnj13*^W53X/+^; t1; gray), after disruption of WT allele (*Kcnj13*^W53X/ΔR^; t2; orange) and the injection of the HD 4X tRNA^Trp.UAG^ to *Kcnj13*^W53X/ΔR^ (t3; purple). **F**, A comparison of the average a-wave and b**-**wave amplitude of *Kcnj13*^W53X/ΔR^ mice after the HD 4X tRNA^Trp.UAG^ at 8 wks and 14 wks. **G**, Comparison of the average c-wave amplitude measured during the experimental time course shows progressive recovery for the mice injected with HD 4X tRNA^Trp.UAG^ at 4, 6, 8, and 14 weeks after treatment.

Lastly, we tested if HD tRNA^Trp.UAG^ could lead to the recovery of the clinical phenotype in an LCA16 *Kcnj13*^W53X/+^ mouse model^23^. After disrupting the WT allele, *Kcnj13*^W53X/□R^, mice were subretinally injected with HD tRNA^Trp.UAG^ (**Fig. 4D**). One of the clinical phenotypes of LCA is severely subnormal or non-detectable Electroretinography (ERG). We performed ERG in mice before and after the disruption of the WT allele to confirm reduced ERG waveforms (a, b, and c-wave amplitude). The a-wave and b-wave amplitudes indicate the function of photoreceptors and inner retinal cells affected by non-functional RPE cells. After treatment of our disease model with HD tRNA^Trp.UAG^ (Fig. 4G), the a-wave amplitude increased from 67.5 ± 14.7 µV to 73.0 ± 14.2 µV (8 wks, P=0.71) and 98.2 ± 9.5 µV (14 wks, P=0.11), which is more than 1.5 times higher. Similarly, the b-wave amplitude increased from 121.8 ± 30.3 µV to 149.9 ± 40.7 µV (8 wks, P=0.39), and 187.0 ± 24.3 µV (14 wks, P=0.05) (Fig. 4F). The c-wave, which purely represents RPE cell function, decreased from 407.03 ± 28.9 µV to 134.9 ± 31.5 µV after the disruption of the WT allele. After treating the mice with HD tRNA^Trp.UAG^, c-wave amplitude increased to 195.9 ± 46.6 µV in four weeks. The c-wave showed an increasing trend despite a slight dip at the 6-week time point, measuring 214.4 ± 29.3 µV at 8 weeks. After 14 weeks, the c-wave amplitude measured 208.5 + 29.9 µV, confirming a durable therapeutic outcome (**Fig. 4F**). Overall, these results confirm the efficacy of the HDAd virus in transducing RPE cells *in vivo* and support the possibility that ACE-tRNA is effective in reading nonsense mutations to restore *in vivo* phenotypes. The significance was assessed using a one-way ANOVA test.

## Discussion

To allow the correct translation of a UGG codon mutated to a UAG premature termination codon, we engineered tRNA-Trp with a CUA anticodon, which is then endogenously charged with tryptophan by tryptophanyl-tRNA synthetase. tRNA^Trp.UAG^ allowed the readthrough of the UAG premature stop codon by incorporating tryptophan during translation, as shown in our GFP reporter expression studies. The number and type of tRNA genes in tryptophan vary across species. In humans, there are at least seven anticodon CCA genes (1-1 to 5-1). CCA 2-1 tRNA^Trp.UAG^ showed optimal read-through, possibly due to the genomic variations that stabilize the ribosomal translation process. Post-transcriptional modifications of engineered tRNA also ensure their stability and accuracy, reinforcing their critical function in maintaining cellular proteostasis^14,24^. Since multiple copies of the tRNA-Trp gene might be needed to meet the translational demand for tryptophan in protein synthesis, we titrated to use 4X tRNA^Trp.UAG^, which enabled normal translation in both LCA16 hiPSC-RPE and mice.

tRNA^Trp.UGA^ was previously shown to mediate the readthrough of UGA stop codons in cultured cells^16,25^. We found that tRNA^Trp.UAG^ could achieve therapeutic rescue of clinical phenotypes *in vitro* and *in vivo*, with higher efficiency than reported previously. We noticed that the rescue efficacy improved in iPSC-RPE compared to HEK cells. One possible explanation is that the endogenous PTC-bearing mRNA is expressed at lower levels than the exogenous reporter or Kir1.7 expression plasmid. This suggests that the rescue of endogenous mRNA carrying a PTC is more predictive than that of model cell lines (e.g., HEK293T). There is evidence that ACE-tRNA-Arg incorporated arginine, the WT amino acid, into the growing polypeptide chain through UGA stop codon and anticodon pairing^15^. We reasoned that the physicochemical properties of the amino acids might impact the efficiency and fidelity of suppression, as tryptophan’s bulky aromatic side chain and arginine’s positively charged side chain interact differently with the ribosome and growing peptide chain^26,27^. These differences highlight the importance of tailoring the tRNA-Trp design to a specific nonsense codon and the surrounding sequence context for an optimal readthrough.

The therapeutic outcome of tRNA^Trp.UAG^ requires stable, long-term expression while minimizing toxicity and immune responses. The helper-dependent adenovirus (HDAds) transduction system provides a robust platform for long-term ACE-tRNA expression in post-mitotic hiPSC-RPE and animal models. For ocular gene delivery, HDAd can package large genes for gene augmentation therapy with limited immunogenicity^28^. Previous studies have demonstrated the superior safety profiles of HDAd, which showed reduced inflammatory responses while still enabling sustained transgene expression^29^ due to episomal persistence without genomic integration. This minimizes insertional mutagenesis risks compared to lipid-based systems and conventional viral vectors^30,31^, particularly for systemic delivery^32-34^. Compared to first-generation adenoviral vectors, HDAds have significantly reduced acute toxicity and inflammatory responses and enhanced safety while providing therapeutic efficacy^35^. In our study, subretinal injection of HDAd viruses offered a precise and effective method to deliver tRNA^Trp.UAG^ to retinal cells, making it a candidate for treating nonsense mutations affecting other organs. Another key strength of our approach is that the HDAd vector with the Trp^UAG^ payload provides a gene-agnostic Trp-UAG variant therapy for other diseases based on non-sense mutations that affect a Trp codon to accelerate the clinical course.

The absence of effective treatments for rare or ultra-rare conditions, such as LCA16, highlights the urgent need for innovative therapeutic approaches targeting the genetic basis of these diseases. We suppressed the PTC caused by the W53X mutation in *KCNJ13* through engineered tRNA^Trp.UAG^ treatment provides a compelling proof-of-concept for using ACE-tRNAs in treating LCA16. Nevertheless, there are several challenges for clinical applications. First, long-term efficacy and safety assessments are necessary to determine whether repeated ACE-tRNA treatments are required to maintain therapeutic levels. Furthermore, it is currently unclear whether suppressor tRNAs disrupt cellular metabolism, compete with natural tRNA pools, or require strict regulation of their levels for therapeutic applications^36^. Reassuringly, high levels of UAG suppressor tRNA expression in transgenic mice showed no noticeable effects on tissue morphology during genetic code expansion experiments^9^. Second, while HDAd vectors were effective in our animal model, optimizing the delivery and expression of ACE-tRNA in larger animal models, such as non-human primates, is required to ensure its translatability to human therapy. Investigating alternative delivery methods, including nanoparticles or newer viral vectors with enhanced retinal targeting, may improve efficacy further.

## Conclusion

In summary, this study demonstrates that tRNA^Trp.UAG^-mediated readthrough of the W53X nonsense mutation in *KCNJ13* restored Kir7.1 protein levels and improved cell function in a human iPSC-RPE model of LCA16. Importantly, it also enhanced retinal function in a mouse model of LCA16. Our findings support ACE-tRNA as a promising therapeutic for suppression of nonsense mutation that offers specificity, adaptability, and reduced off-target effects compared to traditional gene therapies. These results pave the way for ACE-tRNA applications across a broader range of genetic diseases, potentially transforming the treatment landscape for gene disorders by enabling safe, effective, and tailored interventions for patients with nonsense mutations.

## Materials and Methods

### ACE-tRNA

The structure and sequence of the ACE-tRNA-containing plasmid used for flow cytometry have been previously described previously^14^. 4x ACE-tRNA^Trp. The UAG^ insert was synthesized by IDT and cloned by Gibson reaction into the G0619 AAV plasmid to create 4x ACE-tRNA^Trp. UAG^ plasmid. The 1x ACE-tRNA^Trp.UAG^ insert was amplified from the G0619-4xACE-tRNA^Trp.UAG^ using LG213/LG214 primers and cloned downstream of the pCMV:mOrange2-TGA-BgHpA sequence at the SphI restriction site via Gibson reaction. 1x and 4x ACE-tRNA^Trp. The UAG^ plasmids used in Supplemental Fig 1 are described in Supplemental Text 1 and 2. LG196/LG219 primers were used to amplify the 4x ACE-tRNA^Trp.UAG^ The insert was cloned into the G1499_pHDAd plasmid via an in-fusion reaction at the AscI restriction site according to the University of Iowa Viral Vector Core (VVC) pHDAd cloning procedure. Primers used for cloning are listed in Supplementary Table 2. The HDAd virus production was assessed using VVC.

### Flow cytometry

Cultured HEK293T cells (10 cm dishes) at 80% confluency were transiently transfected with 2 µg of sfGFP-TAG-containing plasmid and 3 µg of ACE-tRNA-containing plasmid. 24 hours after the transfection, the cells were washed with 1X DPBS and resuspended in 2 % DPBS + FBS solution before sfGFP quantification by flow cytometry on a Becton Dickinson LSR II instrument and analyzed using LSRFortessa Analyzers.

### Cell Culture and Transfection

Human Embryonic Kidney (HEK293) cells were cultured in Dulbecco’s modified Eagle’s medium (DMEM, GIBCO, 12800-017), 10% fetal bovine serum (FBS, Gibco™ Fetal Bovine Serum, Qualified, Cat. 26140095), 2 mmol/L GlutaMAX, and penicillin-streptomycin 100 U/mL. The cells were maintained in a humidified incubator at 37 °C and 5% CO_2_. For transfection, HEK293 cells were seeded to a 35 mm dish and transfected using PolyJet™ (Signa Gen) with pCMV-sfGFP^WT^, pCMV-sfGFP^TAG^, pCMV-eGFP-Kir7.1^WT^, and pCMV-eGFPKir7.1^W53X^ when 70% confluence was reached. 4x ACE-tRNA^Trp. The UAG^ plasmid was co-transfected with either pCMV-sfGFP^TAG^ or pCMV-eGFPKir7.1^W53X^ for read-through of the nonsense mutation.

### Differentiation and maintenance of iPSC-RPE

The WT-hiPSC-RPE and LCA16-hiPSC lines used in this study were generated using hiPSC lines that were cultured and differentiated into RPE according to a previously described protocol^37-39^. Briefly, hiPSCs were maintained on Matrigel-coated plates in the mTeSR Plus medium (STEMCELL Technologies). For differentiation, cells were harvested using ReLeSR (STEMCELL Technologies) to form embryoid bodies (EBs) and maintained as a suspension culture in mTeSR Plus, which was gradually transitioned to Neural Induction medium (NIM; DMEM: F12; 1% N2 supplement, 1% MEM nonessential amino acids, 1% L-glutamine, and 2 mg/mL heparin) by day 4. On day 7, EBs were plated on laminin-coated culture plates for further differentiation into adherent cultures. On day 16, the 3D neural rosette-like structures were removed, and the medium for the remaining adherent cells was changed to retinal differentiation medium (RDM; DMEM/F12 [3.5:1.5], 2% B27 supplement (without retinoic acid), and 1% antibiotic-antimycotic) for further differentiation. After ∼75 days of differentiation, pigmented RPE cells were purified using magnetically activated cell sorting (MACS) as described by Sharma et al.^40^ and plated on a laminin-coated surface of interest for the intended studies. Matrigel was purchased from WiCell, and all other tissue culture reagents were purchased from Thermo Fisher Scientific.

### Immunocytochemistry

HEK293 cells were transfected with pCMV-eGFP-Kir7.1^WT^, pCMV-eGFPKir7.1^W53X,^ or eGFPKir7.1^W53X^ along with 4x ACE-tRNA^Trp.UAG^ plasmids were seeded onto coverslips. After 48 h of plating, the cells were fixed with 4% paraformaldehyde for 15 min at room temperature, followed by three minutes of PBS washing of WT-hiPSC-RPE, LCA16 hiPSC-RPE cells, and LCA16 hiPSC-RPE transduced with HDAd virus carrying ACE-tRNA^Trp. The UAG^ on the transwells was handled in a similar manner. The cells were permeabilized with 0.5% Triton-X 100 at room temperature for 5 min. Membranes were blocked at room temperature for 2 h in a solution containing 2% BSA and 0.25% TritonX 100. Coverslips were incubated with primary antibodies overnight at 4 °C for Kir7.1 C-12 mouse monoclonal (1:300; sc-398810, Santa Cruz Biotechnology, Santa Cruz, CA) and Na-K-ATPase Recombinant Rabbit Monoclonal Antibody (1:300; ST0533, Invitrogen). The coverslips were washed three times in PBS with Tween-20 for 5 min each and incubated with secondary antibodies, such as donkey anti-mouse AlexaFluor-647 (1:3,000) (AB_141607, Invitrogen) and donkey anti-rabbit AlexaFluor-594 (1:3,000) (AB_141637, Invitrogen). Following incubation, the coverslips were washed thrice with PBS containing Tween-20 for 5 min each and stained with DAPI (1:1,000) for 10 min. Coverslips were mounted, and images were acquired using a Nikon-C2 confocal microscope. Offline analysis was performed using the NIS Elements software (Nikon, Melville, NY, USA).

### Patch-clamp electrophysiology

Whole-cell patch-clamp recordings were performed on transfected HEK293 cells to determine the functional effects of the tRNA^Trp.UAG^ treatment. We also recorded freshly dissociated iPSC-RPE cells to evaluate W53X mutation readthrough after HD tRNA^Trp.UAG^ transduction. The extracellular HEPES-buffered Ringer’s solution (HR) included (in mM) 135 NaCl, 1 mM MgCl_2_, 10 HEPES, 1.8 CaCl_2_, 10 mM glucose, and 5 mM KCl. The solution pH was adjusted to 7.4 using NaOH, and an osmolality of 300 mOsm was confirmed. The pipette solution contained 30 mM KCl, 83 K-gluconate, 5.5 EGTA-KOH, 0.5 CaCl_2_, 4 mM MgCl_2,_ 10 mM HEPES, and 4 mM adenosine triphosphate (ATP). The solution pH was adjusted to 7.2 using KOH, and the osmolarity was 280 mOsm. For experiments using 135 mM Rb+, equimolar Na+ in HR was substituted, the pH was adjusted to 7.4 using RbOH, and the solution osmolarity was confirmed to be 300 mOsm. Voltage-clamp experiments used a ramp protocol from -150 to 50 mV from a holding potential of 0 mV. Channel currents were also elicited using a voltage step protocol from +50 to -150 mV in 50 mV steps from a holding potential of 0 mV for 300 ms. The pipette resistance was 2.8-5.5 MΩ during sealing under GΩ conditions. We used Axopatch200B, Digidata1550, and Clampex11 for data acquisition and analyzed the data using Clampfit 11.2 (Molecular Devices, CA, USA).

### Animals

*Kcnj13*^+/W53X^ mice were bred and maintained at the BRMC, the animal facility of the University of Wisconsin-Madison, in a controlled environment with a temperature of 23 ± 2 °C, humidity ranging from 55-60%, and a 12/12 h light-dark cycle. These animals were used to test whether tRNA^Trp.UAG^ therapy restores channel function following the disruption of the wild-type allele. All animals were handled according to the protocol approved by the UW-Madison Institutional Animal Care and Use Committee (IACUC). For ethical purposes, we minimize the number of animals used whenever possible.

### Anesthesia and Subretinal Injection

After weighing, the animals (n=4) were anesthetized by an intraperitoneal injection of ketamine (80 mg/kg) and xylazine (10 mg/kg) cocktails. A drop of 1% tropicamide was applied to the eyes of the mice (n=5) for pupil dilation, and proparacaine hydrochloride was used for topical anesthesia. Mice were placed on a heating pad to maintain their body temperature. A transcorneal subretinal injection was administered to these mice to deliver the viruses. We used a 34-gauge blunt-end needle attached to a 10 µl Hamilton syringe for injections. First, a puncture was made using a 30G disposable needle on the scleral region through which the 34G needle was inserted to reach the subretinal space. Viral particles (2 µL) were delivered to the subretinal space using an UltraMicroPump3 (WPI, Sarasota, FL, USA) at 20 nl/sec. The bleb was visualized to confirm the success of the injection. Antibiotic ointment was applied to the eyes after the injection to prevent infection due to injury.

### W53X mice and WT allele disruption

*Kcnj13*^W53X/+^ mice were created using CRISPR/Cas9 mediated genome engineering, as previously described^23^because *Kcnj13*^W53X/W53X^ mice could not survive beyond postnatal day 1. For this purpose, an *in vivo* evaluation of the tRNA^Trp.UAG^ readthrough was carried out after disrupting the wild-type allele, as described previously. Here, we disrupted the wild-type allele of *Kcnj13*^W53X/+^ by injecting the virus carrying the cas9 protein along with the sgRNA “TAATGGACATGCGCTGGCGCTGG” that is specific to wild-type allele and from here on termed as “*Kcnj13*^W53X/□R”^. The virus was delivered subretinally and followed up with ERG to check the c-wave amplitude arising from RPE cells.

### HD tRNA^Trp.UAG^ injection

All HDAd viruses used in this study were produced by UI Viral Vector Core (University of Iowa). HDAd viruses used in this study either carried CMV-sfGFP, CMV-sfGFP^TAG,^ or 5’Leader-tRNA^Trp. UAG^ (4X_Trp-tRNA). C57BL6 mice were injected with HDAd viruses to evaluate the ability of tRNATrp. UAG to detect tRNATrp. UAG read through TAG nonsense mutations. For the functional rescue study, *Kcnj13*^W53X/□R^ mice that exhibited decreased c-wave amplitude after disruption of the wild-type allele were injected with the HDAd virus carrying 5’Leader-tRNA^Trp.UAG^

### RPE flat-mount imaging

On day 14, C57Bl6 mice injected with HDAd viruses carrying GFP reporters with TAG nonsense mutations and 4X_Trp-tRNA were sacrificed and flat-mounted to evaluate GFP expression after successful readthrough translation by ACE-tRNA. Briefly, the injected eyes were harvested, and an incision was made at the ora serrata using a 30-gauge needle. The eyes were cut along the edge of the cornea, and the lens was removed. The retina was removed by careful dissection, and the remaining eyecup was incised radially to flatten it. The flattened RPE layer was mounted on a cover glass and imaged with NIS-Elements using a Nikon C2 confocal microscope (Nikon Instruments Inc.).

### Electroretinography

The mice were dark-adapted overnight a day before the ERG procedure. ERG was performed as previously described. A drop of 1% tropicamide was applied to the eyes of the mice for pupil dilation. As mentioned previously, mice were anesthetized by injecting a cocktail of ketamine and xylazine at concentrations of 80 and 10 mg/kg, respectively. The entire procedure was performed in a very dim red light, and the body temperature of the mice was maintained at 37°C by placing them on a heating pad. The topical application of 2.5% hemicellulose ophthalmic solution (Goniovisc, HUB Pharmaceuticals LLC, CA) kept the eyes moist and improved electrical conductivity. A contact electrode was placed on the eyes, and ERG was performed using the Espion E3 system with a Ganzfeld color dome (Diagnosis LLC, Lowell, MA, USA) to ensure uniform illumination of the eyes. The seven-step protocol with sequential increments of flash intensities (0.03 to 30 cd.s/m^2^) for 300 ms with 2 s intervals between the flashes was used for scotopic ERG to measure the a-wave and b-wave. The c-wave was acquired by flashing the eyes with 2.5 and 25 cd.s/m^2^ light intensities and was recorded for 4 s. The acquired waveforms were analyzed using the Espion software V6 (Diagnosis LLC, MA) and Origin 2020 software (OriginLab Corp., MA, USA).

### Real-Time PCR

LCA16 hiPSC-RPE cells transduced with HDAd 1X and 4X tRNA^Trp.UAG^ for two weeks were harvested for DNA isolation. Real-time PCR (absolute quantification) was performed using SYBR Green chemistry and specific primers against the ITR region and the C4HSU gene fragment of the HDAd genome (*Supplementary Table 1*). A standard curve was generated using a dilution series of eight different concentrations of the HDAd virus (R^2^=0.95). The target genes (copies/µL) were quantified by comparing the unknown Ct values with the Ct values of the standard curve.

## Statistical Analysis

Statistical analysis was performed using Origin (version 9.0) with a two-tailed Student’s T-test. ANOVA and Tukey’s post-hoc tests were used for multiple comparisons. Data are expressed as the mean ±SEM, and *p<0*.*05*.

## Data Availability

All the datasets used in this study are available from the corresponding author upon request.

## Acknowledgments

We used BioRender, licensed to UW-Madison and NIH BioArt, to generate the artwork shown in all the figures. This work was supported by the National Institute of Health (grant number R24EY032434). This study was supported in part by the Retina Research Foundation M.D. Matthews Professorship (BRP), Daniel M. Albert Chair in Eye Research (BRP), Sandra Lemke Trout Chair in Eye Research (DMG), and RRF Emmett A. Humble Distinguished Directorship (DMG) of the McPherson Eye Institute, UW-Madison. It was also partially supported by an Unrestricted Grant from Research to Prevent Blindness, Inc. to the UW-Madison Department of Ophthalmology and Visual Sciences. The authors also thank the National Institutes of Health (NIH) of the University of Wisconsin-Madison (P30 EY016665) and S10OD026957.

## Author Contributions

**BRP** and **CAA** conceived the study, designed the experiments, and supervised the research. **EA, PKS, LG, AAS, DS, GMH, MAFZ, SMY**, and **MK** conducted the experiments and collected data. **EA, PKS, LG, MK, CAA**, and **BRP** performed data analysis and contributed to figure generation. **BRP, DMG**, and **CAA** provided the essential resources and funding for the project. **EA, PKS, LG, DS, SMY, DMG, CAA**, and **BRP** contributed to the manuscript writing and revision. **BRP, DMG**, and **CAA** managed the project and coordinated the efforts of all contributors. All authors reviewed and approved the final version of the manuscript.

## References

1 Cartegni, L., Chew, S. L. & Krainer, A. R. Listening to silence and understanding nonsense: exonic mutations that affect splicing. Nat Rev Genet 3, 285–298 (2002). 10.1038/nrg775

2 Mort, M., Ivanov, D., Cooper, D. N. & Chuzhanova, N. A. A meta-analysis of nonsense mutations causing human genetic disease. Hum Mutat 29, 1037–1047 (2008). 10.1002/humu.20763

3 Finkel, R. S. Read-through strategies for suppression of nonsense mutations in Duchenne/ Becker muscular dystrophy: aminoglycosides and ataluren (PTC124). J Child Neurol 25, 1158–1164 (2010). 10.1177/0883073810371129

4 Keeling, K. M., Xue, X., Gunn, G. & Bedwell, D. M. Therapeutics based on stop codon readthrough. Annu Rev Genomics Hum Genet 15, 371–394 (2014). 10.1146/annurev-genom-091212-153527

5 Spelier, S., van Doorn, E. P. M., van der Ent, C. K., Beekman, J. M. & Koppens, M. A. J. Readthrough compounds for nonsense mutations: bridging the translational gap. Trends Mol Med 29, 297–314 (2023). 10.1016/j.molmed.2023.01.004

6 Kantor, A., McClements, M. E. & MacLaren, R. E. CRISPR-Cas9 DNA Base-Editing and Prime-Editing. Int J Mol Sci 21 (2020). 10.3390/ijms21176240

7 Drag, S., Dotiwala, F. & Upadhyay, A. K. Gene Therapy for Retinal Degenerative Diseases: Progress, Challenges, and Future Directions. Invest Ophthalmol Vis Sci 64, 39 (2023). 10.1167/iovs.64.7.39

8 Cho, G. Y., Schaefer, K. A., Bassuk, A. G., Tsang, S. H. & Mahajan, V. B. Crispr Genome Surgery in the Retina in Light of Off-Targeting. Retina 38, 1443–1455 (2018). 10.1097/IAE.0000000000002197

9 Porter, J. J., Heil, C. S. & Lueck, J. D. Therapeutic promise of engineered nonsense suppressor tRNAs. Wiley Interdiscip Rev RNA 12, e1641 (2021). 10.1002/wrna.1641

10 Ward, C. et al. Mechanisms and Delivery of tRNA Therapeutics. Chem Rev 124, 7976–8008 (2024). 10.1021/acs.chemrev.4c00142

11 Coller, J. & Ignatova, Z. tRNA therapeutics for genetic diseases. Nat Rev Drug Discov 23, 108–125 (2024). 10.1038/s41573-023-00829-9

12 Fernandez, I. S. et al. Unusual base pairing during the decoding of a stop codon by the ribosome. Nature 500, 107–110 (2013). 10.1038/nature12302

13 Ko, W., Porter, J. J., Sipple, M. T., Edwards, K. M. & Lueck, J. D. Efficient suppression of endogenous CFTR nonsense mutations using anticodon-engineered transfer RNAs. Mol Ther Nucleic Acids 28, 685–701 (2022). 10.1016/j.omtn.2022.04.033

14 Lueck, J. D. et al. Engineered transfer RNAs for suppression of premature termination codons. Nat Commun 10, 822 (2019). 10.1038/s41467-019-08329-4

15 Pezzini, S. et al. Engineered tRNAs efficiently suppress CDKL5 premature termination codons. Sci Rep 14, 31791 (2024). 10.1038/s41598-024-82766-0

16 Blomquist, V. G. et al. Transfer RNA-mediated restoration of potassium current and electrical correction in premature termination long-QT syndrome hERG mutants. Molecular therapy. Nucleic acids 34, 102032 (2023). 10.1016/j.omtn.2023.102032

17 Wang, J. et al. AAV-delivered suppressor tRNA overcomes a nonsense mutation in mice. Nature 604, 343–348 (2022). 10.1038/s41586-022-04533-3

18 Wang, J., Gao, G. & Wang, D. Developing AAV-delivered nonsense suppressor tRNAs for neurological disorders. Neurotherapeutics 21, e00391 (2024).

19 Kumar, M. & Pattnaik, B. R. Focus on Kir7.1: physiology and channelopathy. Channels (Austin) 8, 488–495 (2014). 10.4161/19336950.2014.959809

20 Pattnaik, B. R. et al. A Novel KCNJ13 Nonsense Mutation and Loss of Kir7.1 Channel Function Causes Leber Congenital Amaurosis (LCA16). Hum Mutat 36, 720–727 (2015). 10.1002/humu.22807

21 Wang, T. Y. et al. A novel viral vaccine platform based on engineered transfer RNA. Emerg Microbes Infect 12, 2157339 (2023). 10.1080/22221751.2022.2157339

22 Alfonzo, J. D., Blanc, V., Estevez, A. M., Rubio, M. A. & Simpson, L. C to U editing of the anticodon of imported mitochondrial tRNA(Trp) allows decoding of the UGA stop codon in Leishmania tarentolae. EMBO J 18, 7056–7062 (1999). 10.1093/emboj/18.24.7056

23 Kabra, M. et al. Nonviral base editing of KCNJ13 mutation preserves vision in a model of inherited retinal channelopathy. J Clin Invest 133 (2023). 10.1172/JCI171356

24 Albers, S. et al. Engineered tRNAs suppress nonsense mutations in cells and in vivo. Nature 618, 842–848 (2023). 10.1038/s41586-023-06133-1

25 Porter, J. J., Ko, W., Sorensen, E. G. & Lueck, J. D. Optimization of ACE-tRNAs function in translation for suppression of nonsense mutations. Nucleic Acids Res 52, 14112–14132 (2024). 10.1093/nar/gkae1112

26 Dougherty, D. A. Cation-pi interactions in chemistry and biology: a new view of benzene, Phe, Tyr, and Trp. Science 271, 163–168 (1996). 10.1126/science.271.5246.163

27 Yau, W. M., Wimley, W. C., Gawrisch, K. & White, S. H. The preference of tryptophan for membrane interfaces. Biochemistry 37, 14713–14718 (1998). 10.1021/bi980809c

28 Morral, N. et al. Administration of helper-dependent adenoviral vectors and sequential delivery of different vector serotype for long-term liver-directed gene transfer in baboons. Proc Natl Acad Sci U S A 96, 12816–12821 (1999). 10.1073/pnas.96.22.12816

29 Leggiero, E. et al. Helper-dependent adenovirus-mediated gene transfer of a secreted LDL receptor/transferrin chimeric protein reduces aortic atherosclerosis in LDL receptor-deficient mice. Gene Ther 26, 121–130 (2019). 10.1038/s41434-019-0061-z

30 Du, L., Dronadula, N., Tanaka, S. & Dichek, D. A. Helper-dependent adenoviral vector achieves prolonged, stable expression of interleukin-10 in rabbit carotid arteries but does not limit early atherogenesis. Hum Gene Ther 22, 959–968 (2011). 10.1089/hum.2010.175

31 Zhao, Z., Anselmo, A. C. & Mitragotri, S. Viral vector-based gene therapies in the clinic. Bioeng Transl Med 7, e10258 (2022). 10.1002/btm2.10258

32 Brunetti-Pierri, N. et al. Efficient, long-term hepatic gene transfer using clinically relevant HDAd doses by balloon occlusion catheter delivery in nonhuman primates. Mol Ther 17, 327–333 (2009). 10.1038/mt.2008.257

33 Kojima, H. et al. NeuroD-betacellulin gene therapy induces islet neogenesis in the liver and reverses diabetes in mice. Nat Med 9, 596–603 (2003). 10.1038/nm867

34 Brown, B. D. et al. Helper-dependent adenoviral vectors mediate therapeutic factor VIII expression for several months with minimal accompanying toxicity in a canine model of severe hemophilia A. Blood 103, 804–810 (2004). 10.1182/blood-2003-05-1426

35 Poutou, J. et al. Safety and antitumor effect of oncolytic and helper-dependent adenoviruses expressing interleukin-12 variants in a hamster pancreatic cancer model. Gene Ther 22, 696–706 (2015). 10.1038/gt.2015.45

36 Kiselev, A. V. et al. Suppression of nonsense mutations in the Dystrophin gene by a suppressor tRNA gene. Mol Biol (Mosk) 36, 43–47 (2002).

37 Shahi, P. K. et al. Gene Augmentation and Readthrough Rescue Channelopathy in an iPSC-RPE Model of Congenital Blindness. Am J Hum Genet 104, 310–318 (2019). 10.1016/j.ajhg.2018.12.019

38 Steyer, B. et al. Scarless Genome Editing of Human Pluripotent Stem Cells via Transient Puromycin Selection. Stem Cell Reports 10, 642–654 (2018). 10.1016/j.stemcr.2017.12.004

39 Sinha, D. et al. Human iPSC Modeling Reveals Mutation-Specific Responses to Gene Therapy in a Genotypically Diverse Dominant Maculopathy. Am. J. Hum. Genet. 107, 278–292 (2020). 10.1016/j.ajhg.2020.06.011

40 Sharma, R., Bose, D., Montford, J., Ortolan, D. & Bharti, K. Triphasic developmentally guided protocol to generate retinal pigment epithelium from induced pluripotent stem cells. STAR Protoc 3, 101582 (2022). 10.1016/j.xpro.2022.101582

