## Supplemental figures and tables for "Preventing vision loss in a mouse model of Leber Congenital Amaurosis by engineered tRNA"

***Correspondence:**

Bikash R. Pattnaik

Supplemental Information

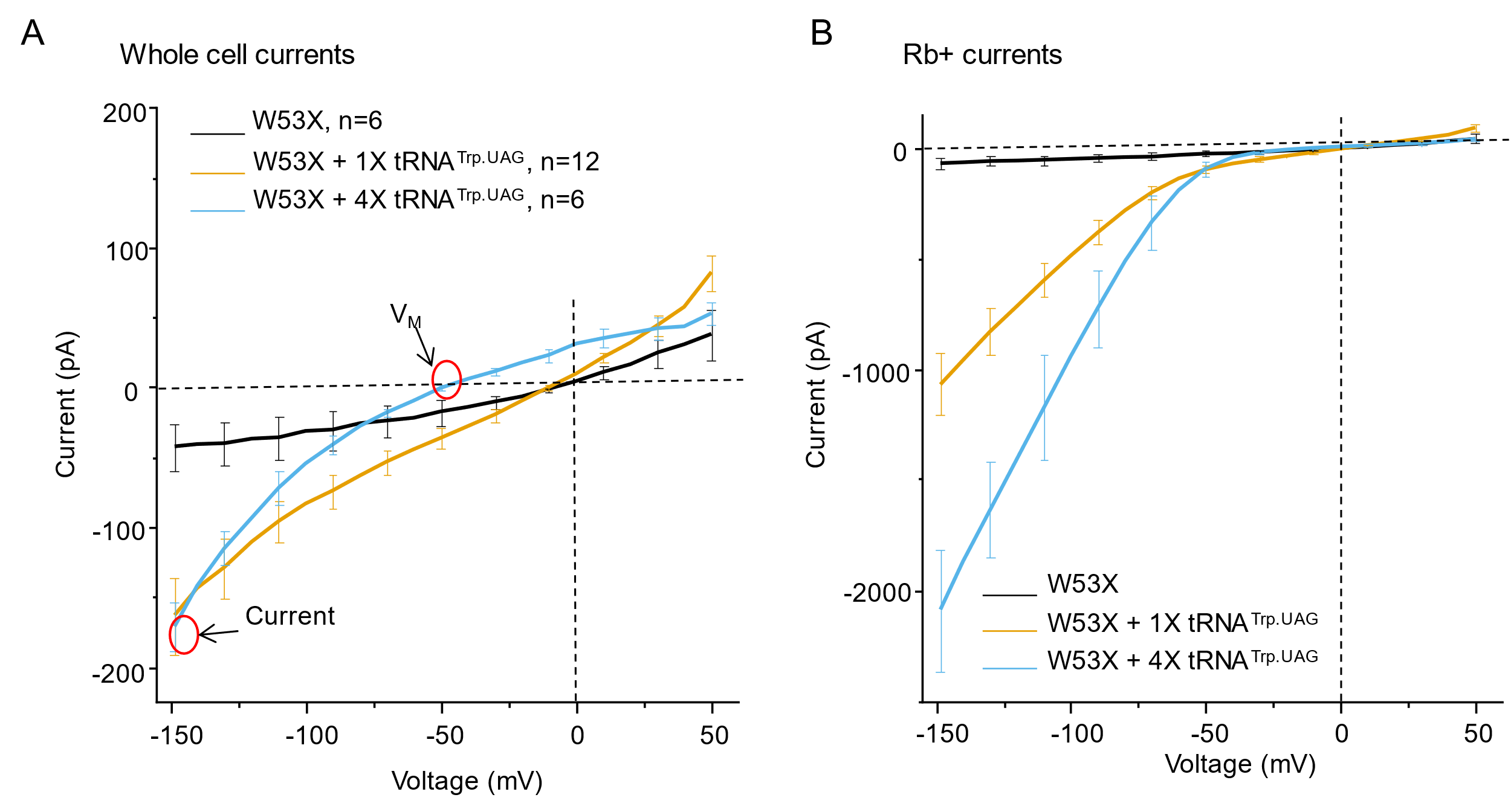

**Supplemental Figure 1. Rescue of K^+^current and membrane potential by only 4X tRNA^Trp.UAG^. (A)** Kir current measured as whole cell current for physiological extracellular 5 mM K+ and **(B)** extracellular 140 mM Rb+ ions for Kir7.1^W53X^ (black line), Kir7.1^W53X^ + **1X** tRNA^Trp.UAG^ (orange line), and Kir7.1^W53X^ + **4X** tRNA^Trp.UAG^ (blue line) channels. As noted in A (red circles), only 4X tRNA^Trp.UAG^ restored both the membrane potential (VM) and the inwardly rectified K+ currents. Data are presented as average values ± SEM.

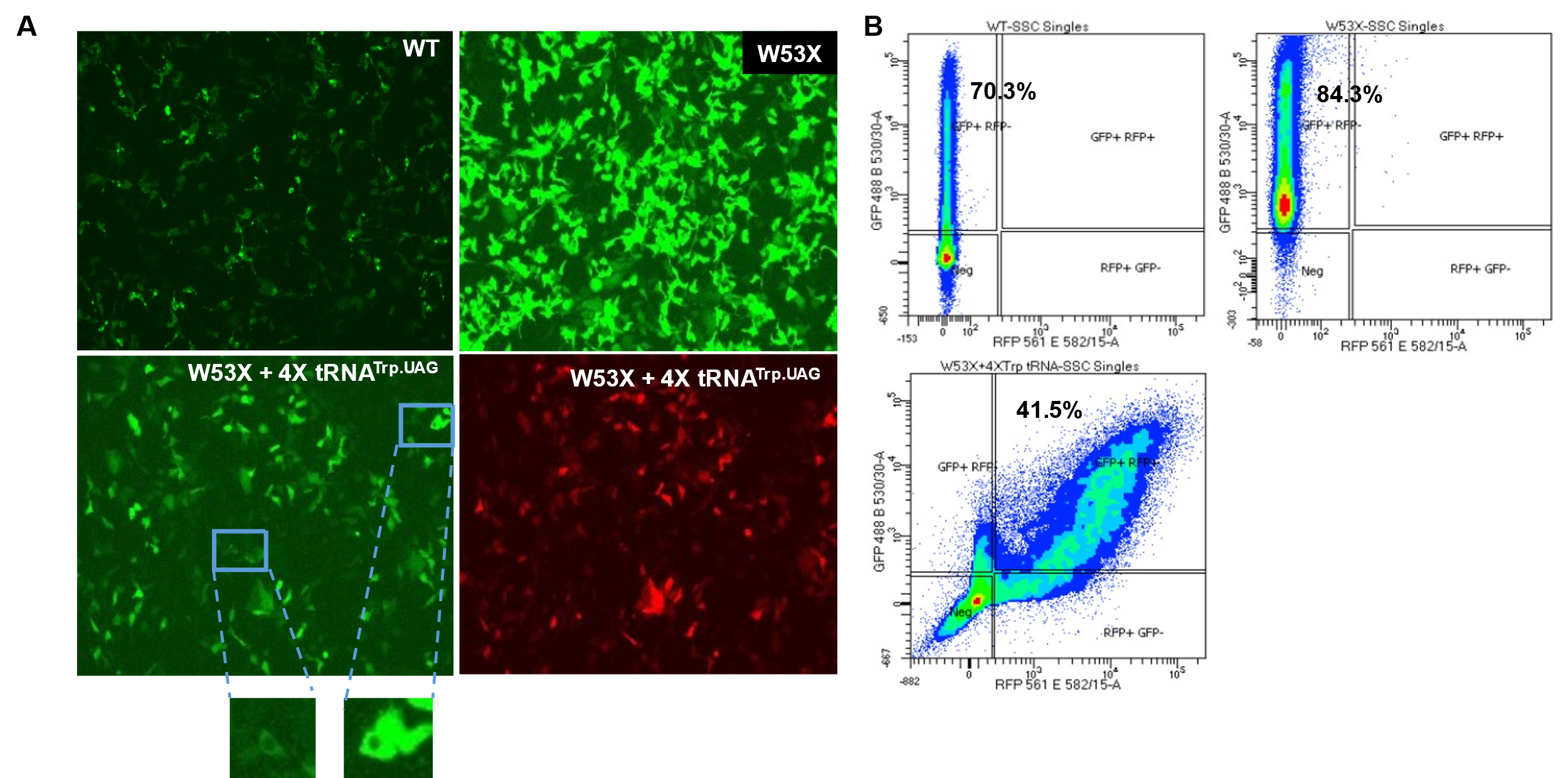

**Supplemental Figure 2. Co-expression of GFP-Kir7.1^W53X^ and tRNA^Trp.UAG^ in HEK293 cells.** **(A)** GFP expression in WT Kir7.1, W53X, and W53X cells treated with tRNA^Trp.UAG^. The red fluorescence in the lower right panel represents cells that express only tRNA^Trp.UAG^, as this plasmid also carries a tdTomato reporter. The cartoon representation is shown in Fig. 2A. **(B)** Flow cytometry demonstrates that the percentage of cells showing GFP expression in wild-type Kir7.1-and W53X mutant-transfected cells was similar, while for co-expression of W53X and 4X tRNA^Trp.UAG, the^ fluorescence distribution shifted to the right, indicating more than 40% co-expression. Scale bars, 50 μm.

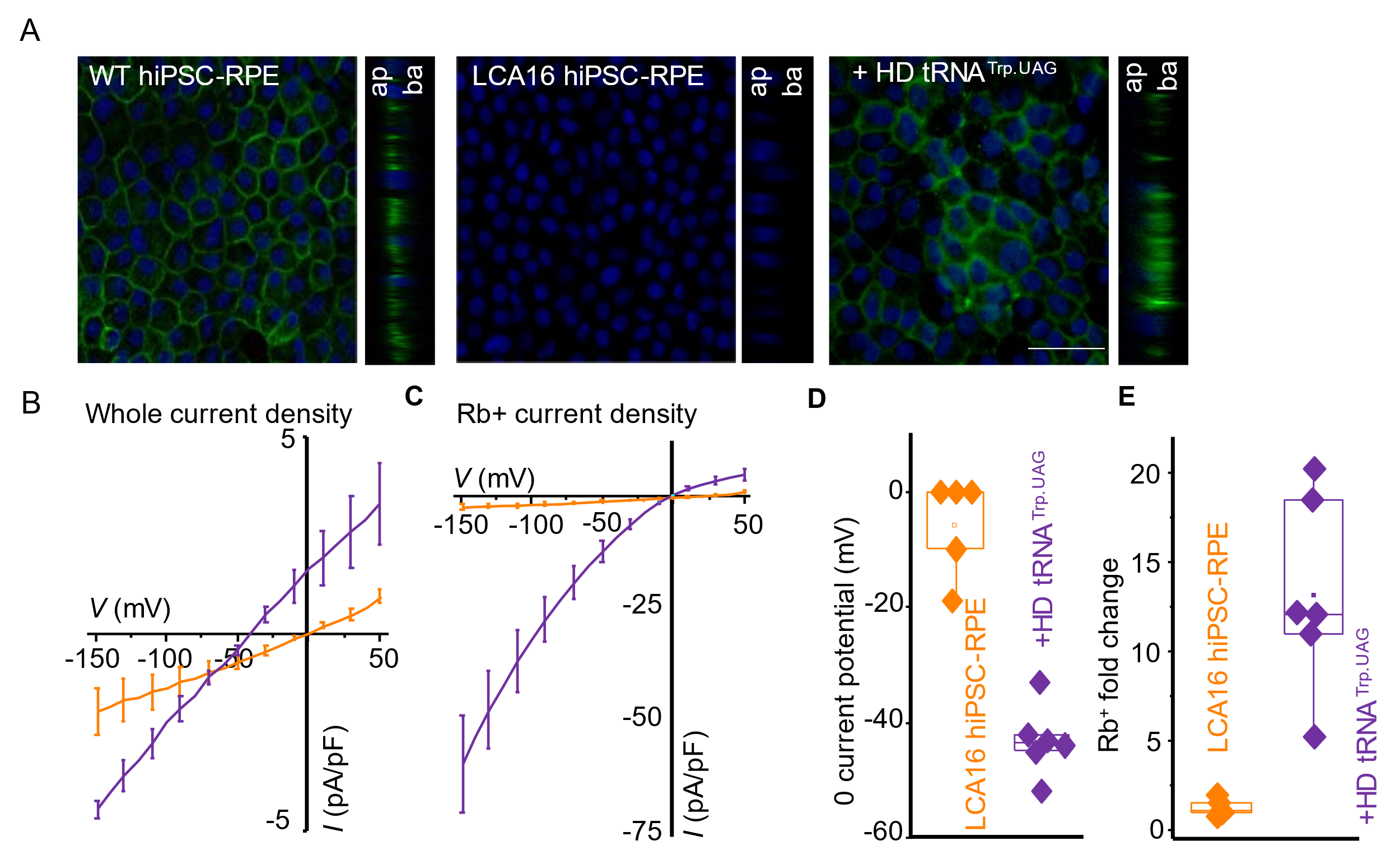

**Supplemental Figure 3. 1X tRNA^Trp.UAG^ treatment of LCA16 hiPSC-RPE cells. (A)** Confocal images of WT hiPSC-RPE, LCA16 hiPSC-RPE, and LCA16 hiPSC-RPE transduced with HD tRNA^Trp.UAG^ shows immunofluorescence localization of Kir7.1 (green) and DAPI staining (blue). Both the apical (ap) and basal (ba) sides are marked in the orthogonal projection. Scale bar, 20 μm; **(B)** Kir current measured as the whole-cell current and **(C)** Rb+ ion current for LCA16 hiPSC-RPE (orange line) and HD tRNATrp. UAG-treated (purple line) cells. The distribution of data points and significance were measured as *P<0.05, **P<0.01, and ***P<0.001 when LCA16 hiPSC-RPE was compared to tRNATrp. UAG-treated groups using two-tailed unpaired Student's t-test. **(D)** Resting membrane potentials for individual cells showing LCA16 hiPSC-RPE (orange quadrangles) and LCA16 hiPSC-RPE cells treated with HD tRNA^Trp.UAG^ (purple quadrangles). **(E)** The inward current-fold increase by extracellular Rb+ was measured at -150 mV for individual cells, as in D.

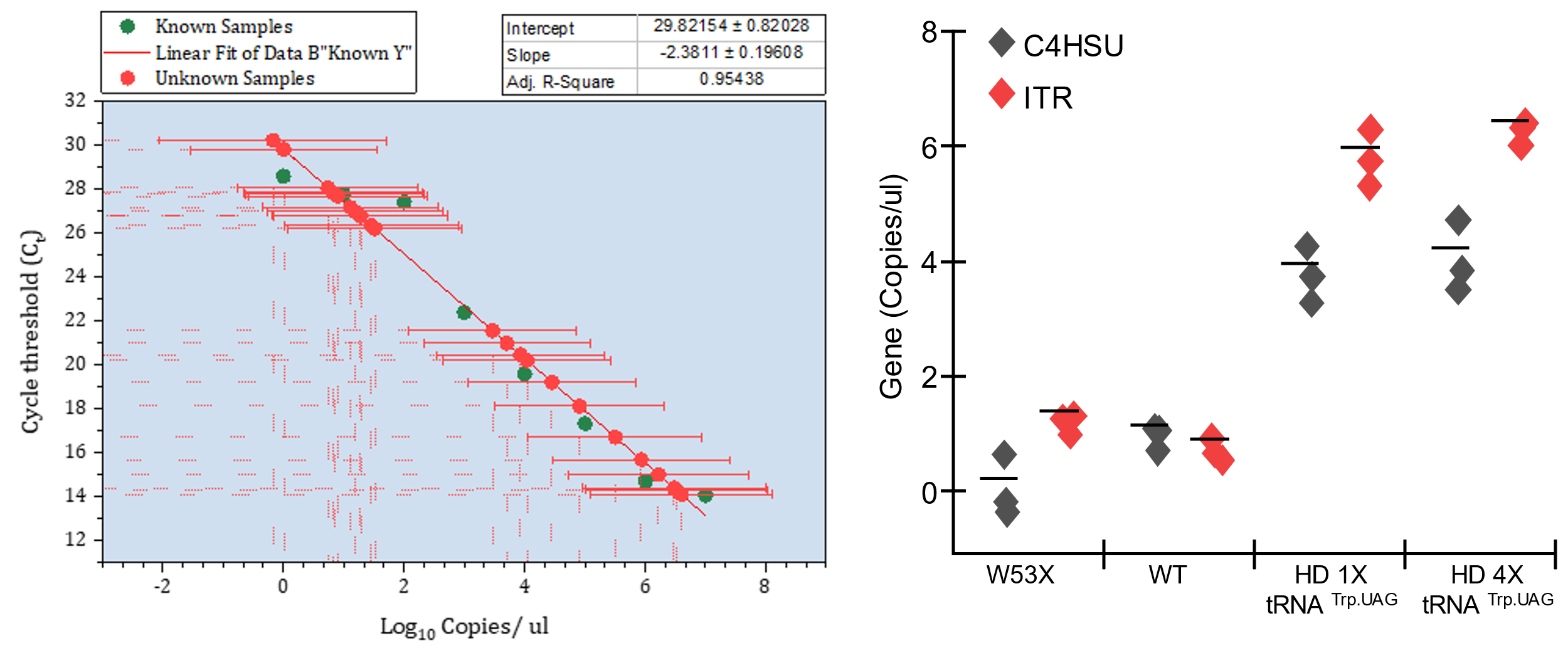

**Supplemental Figure 4.** LCA16 hiPSC-RPE cells were transduced with HD tRNA^Trp.UAG^ 1X and 4X. Absolute quantification of the HD viral genome (ITR and C4HSU, copies/µL) by real-time PCR (n=3, biological controls).

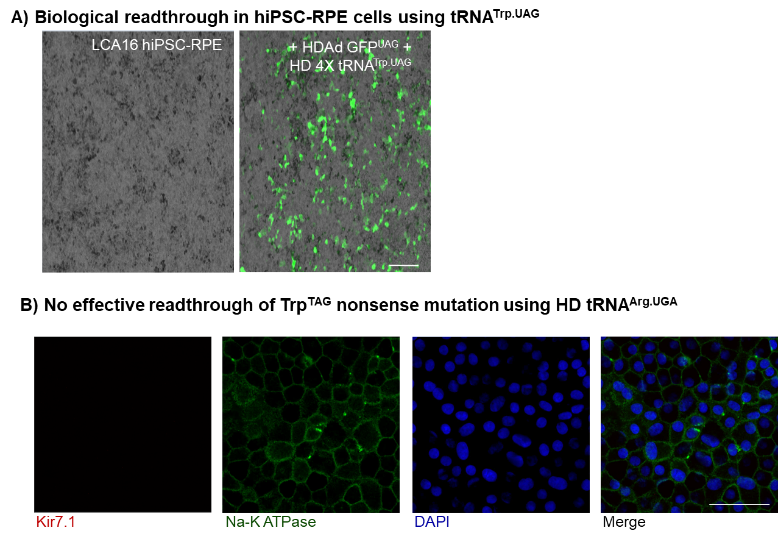

**Supplemental Figure 5. ACE-tRNA readthrough is specific to Ace-tRNA^Arg. TGA^ did not read through the Trp-TAG nonsense codon.** (A) GFP expression was observed as GFP readthrough after transduction of LCA16 hiPSC-RPE cells with CMV_sfGFP^TAG^ and HD tRNA^Trp.^**^UAG^**. (B) Specificity of HD tRNA^Trp.UAG^ was confirmed by the absence of Kir7.1 readthrough in LCA16 hiPSC-RPE cells after transduction with HD tRNA^Arg.^**^UGA^**.

**Supplemental Text 1** Payload_mOrange2-TAG_1xtRNA_Trp_TAG

CTGCGCGCTCGCTCGCTCACTGAGGCCGCCCGGGCAAAGCCCGGGCGTCGGGCGACCTTTGGTCGCCCGGCCTCAGTGAGCGAGCGAGCGCGCAGAGAGGGAGTGGAATTCACGCGTGGATCTGAATTCAATTCACGCGTGGTACCCAATTGAGATCTTCTAGAGTTTAAACCTCGAGAAGCTTTTAATTAAGCTAGCGGATAAGCTTGGGAGTTCCGCGTTACATAACTTACGGTAAATGGCCCGCCTGGCTGACCGCCCAACGACCCCCGCCCATTGACGTCAATAATGACGTATGTTCCCATAGTAACGCCAATAGGGACTTTCCATTGACGTCAATGGGTGGAGTATTTACGGTAAACTGCCCACTTGGCAGTACATCAAGTGTATCATATGCCAAGTACGCCCCCTATTGACGTCAATGACGGTAAATGGCCCGCCTGGCATTATGCCCAGTACATGACCTTATGGGACTTTCCTACTTGGCAGTACATCTACGTATTAGTCATCGCTATTACCATGGTGATGCGGTTTTGGCAGTACATCAATGGGCGTGGATAGCGGTTTGACTCACGGGGATTTCCAAGTCTCCACCCCATTGACGTCAATGGGAGTTTGTTTTGGCACCAAAATCAACGGGACTTTCCAAAATGTCGTAACAACTCCGCCCCATTGACGCAAATGGGCGGTAGGCGTGTACGGTGGGAGGTCTATATAAGCAGAGCTGGTTTAGTGAACCGTCAGATCCGCTAGCGCCACCATGGTGAGCAAGGGCGAGGAGAATAACATGGCCATCATCAAGGAGTTCATGCGCTTCAAGGTGCGCATGGAGGGCTCCGTGAACGGCCACGAGTTCGAGATCGAGGGCGAGGGCGAGGGCCGCCCCTACGAGGGCTTTCAGACCGCTAAGCTGAAGGTGACCAAGGGTGGCCCCCTGCCCTTCGCCTGGGACATCCTGTCCCCTCATTTCACCTACGGCTCCAAGGCCTACGTGAAGCACCCCGCCGACATCCCCGACTACTTCAAGCTGTCCTTCCCCGAGGGCTTCAAGTGGGAGCGCGTGATGAACTACGAGGACGGCGGCGTGGTGACCGTGACCCAGGACTCCTCCCTGCAGGACGGCGAGTTCATCTACAAGGTGAAGCTGCGCGGCACCAACTTCCCCTCCGACGGCCCCGTGATGCAGAAGAAGACCATGGGCTGGGAGGCCTCCTCCGAGCGGATGtagCCCGAGGACGGTGCCCTGAAGGGCAAGATCAAGATGAGGCTGAAGCTGAAGGACGGCGGCCACTACACCTCCGAGGTCAAGACCACCTACAAGGCCAAGAAGCCCGTGCAGCTGCCCGGCGCCTACATCGTCGACATCAAGTTGGACATCACCTCCCACAACGAGGACTACACCATCGTGGAACAGTACGAACGCGCCGAGGGCCGCCACTCCACCGGCGGCATGGACGAGCTGTACAAGTGATAATAGGATCCGATATCACTAGTGCGGCCGCGTCGACTAGAGCTCGCTGATCAGCCTCGACTGTGCCTTCTAGTTGCCAGCCATCTGTTGTTTGCCCCTCCCCCGTGCCTTCCTTGACCCTGGAAGGTGCCACTCCCACTGTCCTTTCCTAATAAAATGAGGAAATTGCATCGCATTGTCTGAGTAGGTGTCATTCTATTCTGGGGGGTGGGGTGGGGCAGGACAGCAAGGGGGAGGATTGGGAAGACAATAGCAGGCATGCTGGGGA*AGCGCTCCGGTTTTTCTGTGCTGAACCTCAGGGGACGCCGACACACGTACACGTC***GACCTCGTGGCGCAATGGTAGCGCGTCTGACTctaGATCAGAAGGtTGCGTGTTCAAGTCACGTCGGGGTCA**GTCCTTTTTTTGCTGGGGAGAGATCGATCTGAGGAACCCCTAGTGATGGAGTTGGCCACTCCCTCTCTGCGCGCTCGCTCGCTCACTGAGGCCGGGCGACCAAAGGTCGCCCGACGCCCGGGCTTTGCCCGGGCGGCCTCAGTGAGCGAGCGAGCGCGCAGAGAGGGAGTGGCC

ITR, CMV promoter, mOrange2-Y156TAG, BgH pA

tRNA:

*5’ leader*, **tRNA Trp UAG**, and terminator

**Supplementary Text 2** -Payload_tdTomatoWT_4xtRNA_Trp_TAG

CCTGCAGGCAGCTGCGCGCTCGCTCGCTCACTGAGGCCGCCCGGGCAAAGCCCGGGCGTCGGGCGACCTTTGGTCGCCCGGCCTCAGTGAGCGAGCGAGCGCGCAGAGAGGGAGTGGCCAACTCCATCACTAGGGGTTCCTTGTAGTTAATGATTAACCCGCCATGCTACTTATCTACGTAGCCATGCTCTAGAGCGGCCGCACGCGTACTAGTAAGGGCCTCGAGGTCGACAGAAGCAGGCTCGGTAACATACGGTCTAGCTATCTGACTAGCGCTCCGGTTTTTCTGTGCTGAACCTCAGGGGACGCCGACACACGTACACGTC**GACCTCGTGGCGCAATGGTAGCGCGTCTGACTctaGATCAGAAGGtTGCGTGTTCAAGTCACGTCGGGGTCA**GTCCTTTTTTTGACAAAGTACTGGTTCCCTTTCCAGCGGGATGCTTTATCTAAACGCAATGAGAGAGGTATTCCTCAGGCCACATCGCTTCCTAGTTCCGCTGGGATCCATCGTTGGCGGCCGAAGCCGCCATTCCATAGTGAGTTCTTCGTCTGTGTCATTCTGTGCCAGATCGTCTGGCAAATAGCCGATCCAGTTTATCTCTCGAAACTATAGTCGTACAGATCGAAATCTTAAGTCAAATCACGCGACTAGACTCAGCTCTATTTTAGTGGTCATGGGTTTTGGTCCCCCCGAGCGGTGCAACCGATTAGGACCATGTAGAACATTAGTTATAAGTCTTCTTTTAAACACAATCTTCCTGCTCAGTGGTACATGGTTATCGTTATTGCTAGCCAGCCTGATAAGTAACAGCGCTCCGGTTTTTCTGTGCTGAACCTCAGGGGACGCCGACACACGTACACGTC**GACCTCGTGGCGCAATGGTAGCGCGTCTGACTctaGATCAGAAGGtTGCGTGTTCAAGTCACGTCGGGGTCA**GTCCTTTTTTTGTATCACTTTTGCCTCAACGACTGCTGCTTTCGCTGTAACCCAAGACAGACAACAGTAACCGCCTTTTGAAGGCAAGTCCTCCGCCTGTGACTAACTGTGCCAAATCGTCTTCCAAACCCCTTATCCAATTTAACTCACCAAATTATTGCGATACAGACCCTAATTTCACATCATATGACACTAATTGCCTCTGCCAAAATTCTGTCCTCAAGCGTTTTAGTTCGCCCCAGTAATGTTGCCAATAAGGACCACCAAATCCGCATGTTACAGGACTTCTTATAAATTCTTTTTTCGTGGGGAGCAGCGGATCTTAATGGATGGCGCCAGCTGGTATGGAAGCTAATAGCGCCGGTGAGAGGGTAATCAGCCGTCTCCACCAACACAACGCTATCGGGTCATAAGCGCTCCGGTTTTTCTGTGCTGAACCTCAGGGGACGCCGACACACGTACACGTC**GACCTCGTGGCGCAATGGTAGCGCGTCTGACTctaGATCAGAAGGtTGCGTGTTCAAGTCACGTCGGGGTCA**GTCCTTTTTTTGTATAGTGCGCTGATCGGAGACGAATTAAAAACACGAGTTCCCAAAACCAGGCGGGCTCGCCACGTCGGCTAATCCTGGTACATTATGTGAACAATGTTCTGAAGAAAATTTGTGAAAGAAGGACGGGTCATCGCCTACTATTAGCAACAACGGTCGGCCACACCTTCCATTGTCGTGGCCACGCTCGGATTACACGGCAGAGGTGCTTGTGTTCCGACAGGCTAGCATATTATCCTAAGGCGTTACCCCAATCGTTTACCGTCGGATTTGCTATAGCCCCTGAACGCTACATGTACGAAACCATGTTATGTATGCACTAGGTCAACAATAGGACATAGCCTTGTAGTTAACACGTAGCCCGGTCGTATAAGTACAGTAGACCCTTCGCCGGCATCCTATTAGCGCTCCGGTTTTTCTGTGCTGAACCTCAGGGGACGCCGACACACGTACACGTC**GACCTCGTGGCGCAATGGTAGCGCGTCTGACTctaGATCAGAAGGtTGCGTGTTCAAGTCACGTCGGGGTCA**GTCCTTTTTTTGTACTGGTCTGCGGAGCACTCTGGTTATGCATATGGTCCACAGGAGAATTCTCGACTTATTAATAGTAATCAATTACGGGGTCATTAGTTCATAGCCCATATATGGAGTTCCGCGTTACATAACTTACGGTAAATGGCCCGCCTGGCTGACCGCCCAACGACCCCCGCCCATTGACGTCAATAATGACGTATGTTCCCATAGTAACGCCAATAGGGACTTTCCATTGACGTCAATGGGTGGAGTATTTACGGTAAACTGCCCACTTGGCAGTACATCAAGTGTATCATATGCCAAGTACGCCCCCTATTGACGTCAATGACGGTAAATGGCCCGCCTGGCATTATGCCCAGTACATGACCTTATGGGACTTTCCTACTTGGCAGTACATCTACGTATTAGTCATCGCTATTACCATGGTGATGCGGTTTTGGCAGTACATCAATGGGCGTGGATAGCGGTTTGACTCACGGGGATTTCCAAGTCTCCACCCCATTGACGTCAATGGGAGTTTGTTTTGGCACCAAAATCAACGGGACTTTCCAAAATGTCGTAACAACTCCGCCCCATTGACGCAAATGGGCGGTAGGCGTGTACGGTGGGAGGTCTATATAAGCAGAGCTGGTTTAGTGAACCGTCAGATCCGCTAGCGCCACCATGGTGAGCAAGGGCGAGGAGGTCATCAAAGAGTTCATGCGCTTCAAGGTGCGCATGGAGGGCTCCATGAACGGCCACGAGTTCGAGATCGAGGGCGAGGGCGAGGGCCGCCCCTACGAGGGCACCCAGACCGCCAAGCTGAAGGTGACCAAGGGCGGCCCCCTGCCCTTCGCCTGGGACATCCTGTCCCCCCAGTTCATGTACGGCTCCAAGGCGTACGTGAAGCACCCCGCCGACATCCCCGATTACAAGAAGCTGTCCTTCCCCGAGGGCTTCAAGTGGGAGCGCGTGATGAACTTCGAGGACGGCGGTCTGGTGACCGTGACCCAGGACTCCTCCCTGCAGGACGGCACGCTGATCTACAAGGTGAAGATGCGCGGCACCAACTTCCCCCCCGACGGCCCCGTAATGCAGAAGAAGACCATGGGCTGGGAGGCCTCCACCGAGCGCCTGTACCCCCGCGACGGCGTGCTGAAGGGCGAGATCCACCAGGCCCTGAAGCTGAAGGACGGCGGCCACTACCTGGTGGAGTTCAAGACCATCTACATGGCCAAGAAGCCCGTGCAACTGCCCGGCTACTACTACGTGGACACCAAGCTGGACATCACCTCCCACAACGAGGACTACACCATCGTGGAACAGTACGAGCGCTCCGAGGGCCGCCACCACCTGTTCCTGGGGCATGGCACCGGCAGCACCGGCAGCGGCAGCTCCGGCACCGCCTCCTCCGAGGACAACAACATGGCCGTCATCAAAGAGTTCATGCGCTTCAAGGTGCGCATGGAGGGCTCCATGAACGGCCACGAGTTCGAGATCGAGGGCGAGGGCGAGGGCCGCCCCTACGAGGGCACCCAGACCGCCAAGCTGAAGGTGACCAAGGGCGGCCCCCTGCCCTTCGCCTGGGACATCCTGTCCCCCCAGTTCATGTACGGCTCCAAGGCGTACGTGAAGCACCCCGCCGACATCCCCGATTACAAGAAGCTGTCCTTCCCCGAGGGCTTCAAGTGGGAGCGCGTGATGAACTTCGAGGACGGCGGTCTGGTGACCGTGACCCAGGACTCCTCCCTGCAGGACGGCACGCTGATCTACAAGGTGAAGATGCGCGGCACCAACTTCCCCCCCGACGGCCCCGTAATGCAGAAGAAGACCATGGGCTGGGAGGCCTCCACCGAGCGCCTGTACCCCCGCGACGGCGTGCTGAAGGGCGAGATCCACCAGGCCCTGAAGCTGAAGGACGGCGGCCACTACCTGGTGGAGTTCAAGACCATCTACATGGCCAAGAAGCCCGTGCAACTGCCCGGCTACTACTACGTGGACACCAAGCTGGACATCACCTCCCACAACGAGGACTACACCATCGTGGAACAGTACGAGCGCTCCGAGGGCCGCCACCACCTGTTCCTGTACGGCATGGACGAGCTGTACAAGTGACAATTCGCCCTATAGTGAGTCGTATTACGCGCGCAGCGGCCGACCATGGCCCAACTTGTTTATTGCAGCTTATAATGGTTACAAATAAAGCAATAGCATCACAAATTTCACAAATAAAGCATTTTTTTCACTGCATTCTAGTTGTGGTTTGTCCAAACTCATCAATGTATCTTATCATGTCTGGATCTCCGGACACGTGCGGACCGAGCGGCCGCTCTAGAGCATGGCTACGTAGATAAGTAGCATGGCGGGTTAATCATTAACTACAAGGAACCCCTAGTGATGGAGTTGGCCACTCCCTCTCTGCGCGCTCGCTCGCTCACTGAGGCCGGGCGACCAAAGGTCGCCCGACGCCCGGGCGGCCTCAGTGAGCGAGCGAGCGCGCAGCTGCCTGCAGG

ITR, CMV promoter, TdTomato WT, SV40 pA

tRNA:

*5’ leader*, **tRNA Trp UAG**, and terminator

**Supplemental Table 1**: Primers used for absolute quantification of ITR and C4HSU

| **Primer name** | **Primer sequence (5’ – 3’)** |
| --- | --- |
| ITR_qRT Forward primer | GGGGGTGGAGTTTGTGACG |
| ITR_qRT Reverse primer | GTCACCCGCCCCGTT |
| C4HSU_qRT Forward primer | CCCCGCTACCCCAATCC |
| C4HSU_qRT Reverse primer | TTAGCTTTTTTGGGTGATTTTTCC |

**Supplemental Table 2**: Primers used for cloning

| Primer name | Primer sequence (5’ – 3’) |
| --- | --- |
| LG213 | GATTGGGAAGACAATAGCAGGCATGCTGGGGAAGCGCTCCGGTTTTTCTGTGCTG |
| LG214 | GTTCCTCAGATCGATCTCTCCCCAGCAAAAAAAGGACTGACCCCGACGT |
| LG196 | TTGGCGCGCCGATAAGCTTGGGAGTTCCGCGTTACATAACTTACG |
| LG219 | TTGGCGCGCCCAAAAAAAGGACTGACCCCGACGTG |
